# Modeling the development of cortical responses in primate dorsal (“where”) pathway to optic flow using hierarchical neural field models

**DOI:** 10.1101/2022.08.18.504366

**Authors:** Anila Gundavarapu, V Srinivasa Chakravarthy

**Affiliations:** Computational Neuroscience Lab, Indian Institute of Technology Madras, Chennai, India, 600036; Center for Complex Systems and Dynamics, Indian Institute of Technology Madras, Chennai, India, 600036

## Abstract

Although there is a plethora of modelling literature dedicated to the object recognition processes of the ventral (“what”) pathway of primate visual systems, modelling studies on the motion sensitive regions like the Medial superior temporal area (MST) of the dorsal (“where”) pathway are relatively scarce. Neurons in the MST area of the macaque monkey respond selectively to different types of optic flow sequences such as radial and rotational flows. We present three models that are designed to simulate the computation of optic flow performed by the MST neurons in primates. The first two models are each composed of 3 stages: the first stage comprises the Direction Selective Mosaic Network (DSMN), the second stage comprises the Cell Plane Network (CPNW) or the Hebbian Network (HBNW) and the third stage comprises the optic flow network (OF). The three stages roughly correspond to V1-MT-MST areas respectively in the primate motion pathway. Both these models are trained stage by stage using a biologically plausible variation of Hebbian learning. On the other hand, model-3 consists of the Velocity Selective Mosaic Network (VSMN) followed by a convolutional neural network (CNN) which is trained using supervised backpropagation algorithm. We created various dot configurations that can move in translational, radial, and rotational trajectories to make training and test set. The simulation results show that, while neurons in model-1 and model-2 could account for MSTd cell properties found neurobiologically, model-3 neuron responses are consistent with the idea of functional hierarchy in the macaque motion pathway. These results also suggest that the deep learning models could offer a computationally elegant and biologically plausible solution to simulate the development of cortical responses of the primate motion pathway.

## 1. INTRODUCTION

Optic flow refers to global motion in the retinal image caused by the motion of the observer relative to the world (Gibson, 1950). It is used to compute useful quantities such as heading direction, which specifies the direction of self-motion relative to direction of gaze, translational and rotational velocity of the observer. The Middle Temporal (MT) area contains many direction-selective cells (Maunsell & van Essen, 1983a, 1983b; Rodman & Albright, 1987) that encode the flow field of the evolving retinal image (Britten et al., 1993; Bülthoff et al., 1989; Lappe et al., 1996; J. Movshon et al., 1992; Newsome et al., 1990; Wang et al., 1989). In Medial Superior Temporal (MST) area, many neurons respond to spatially extended random dot optic flow patterns (Duffy & Wurtz, 1991b; Saito et al., 1986; Tanaka & Saito, 1989). Cells in MST area have large receptive fields ∼15° - 100° (Duffy & Wurtz, 1991a, 1991b), that respond selectively to expansion, rotation and combination motion stimuli that are generated due to observer motion (Graziano et al., 1994; Saito et al., 1986; Tanaka & Saito, 1989). MST cells receive their primary input from MT (Boussaoud et al., 1990; Desimone & Ungerleider, 1986) where the initial processing of optic flow involves computation of direction and speed within a small region of the visual field. The emergence of MST and MT cell responses poses an important question: “How can local MT motion estimates be organized into the global selectivity for optic flow that helps in estimation of heading?” Various models have been proposed to elucidate the possible implementation of optic flow and heading estimation in the area MST.

### Related modelling studies

Smith and colleagues (Smith et al., 2006) proposed a computational model in which optic flow selectivity is derived by integrating over MT region where neurons are selective for local direction of optic flow at each point in the receptive field. However, it does not address how MST cells may facilitate navigation by helping to compute estimates of heading. Lappe and Rauschecker (Lappe & Rauschecker, 1993) devised a network model of heading estimation in which a population of neurons code for specific heading direction. The line of modeling proposed by Perrone and colleagues (Perrone, 1992; Perrone & Stone, 1994; Stone & Perrone, 1997) took a different view where individual units directly code for heading direction as an early step in the cascade of processing necessary for self-motion perception and navigation. Some authors proposed that there are three main classes of biological models of neural processing at MST: differential motion, decomposition and template models (Browning et al., 2008).

These findings provide compelling evidence that heading perception is based on the evolution of the optic flow field but are not especially informative about the nature of the underlying neural mechanisms such as temporal dynamics that arise due to the integration of information over time. However, the neural network model for the extraction of optic flow proposed by Fukushima and colleague (Fukushima, 2008; Tohyama & Fukushima, 2005) suggests a different approach: the vector field hypothesis that any flow field can be mathematically decomposed into elementary flow components such as divergence and curl.

### Recent deep learning approaches of modeling neuron selectivity in visual system

Biological realism is the primary concern of most of the neuroscience models (Hubel & Wiesel, 1962; Pack & Born, 2008; Rousselet et al., 2002; Rust et al., 2006; Serre et al., 2007; Simoncelli & Heeger, 1998). These models are basically designed to account for anatomical and neurophysiological data and did not scale up to solving real world tasks. Currently feed-forward convolutional neural networks (CNNs) (LeCun et al., 2015), are the state-of-the-art for object classification tasks such as ImageNet, on occasions surpassing human performance (He et al., 2015). Recently several studies (Agrawal et al., 2014; Güçlü & van Gerven, 2015; Kriegeskorte, 2015) have begun to assess the convolutional neural network as a model for biological vision systems by comparing the internal representations and performance levels between artificial and biologically realistic neural network models. Studies also showed (Kriegeskorte, 2015) that deep convolutional neural networks trained on object recognition tasks not only have architectural similarities but also learn representations that are similar to the representations of the neurons in the ventral pathway. However, their suitability and performance on the dorsal-stream regions is an open area of research.

In this paper we describe: (i) a competitive learning algorithm/design (model-2) that shows emergence of optic flow sensitivity, (ii) a deep neural architecture (model-3) that can learn motion-related properties similar to the representations of neurons in MT/MST. Section 2 (Model architecture and learning rules) describes the architecture of the three models. Even though we described it as three distinct models, they have some common components; components that differ between the models perform equivalent functions (Fig. 1). The main structure of these models is a multi-stage neural network composed of the initial stage simulating the direction selective neurons of V1, the middle stage simulating translational motion selective neurons of MT and the output stage simulating the optic flow selective MST neurons. An important common feature of all these proposed models is the presence of 2D layers of neurons with lateral connections trained by asymmetric Hebbian learning (stage-1). The combination of lateral connectivity and asymmetric Hebbian learning provides an opportunity for extracting motion information. Section 2 also discusses the training procedure of the three network models. Section 3 shows the simulation results and compares the performances of the model with the biological properties of the MST neurons. Section 4 and Section 5 consist of discussion and conclusion respectively.

**Figure 1:**
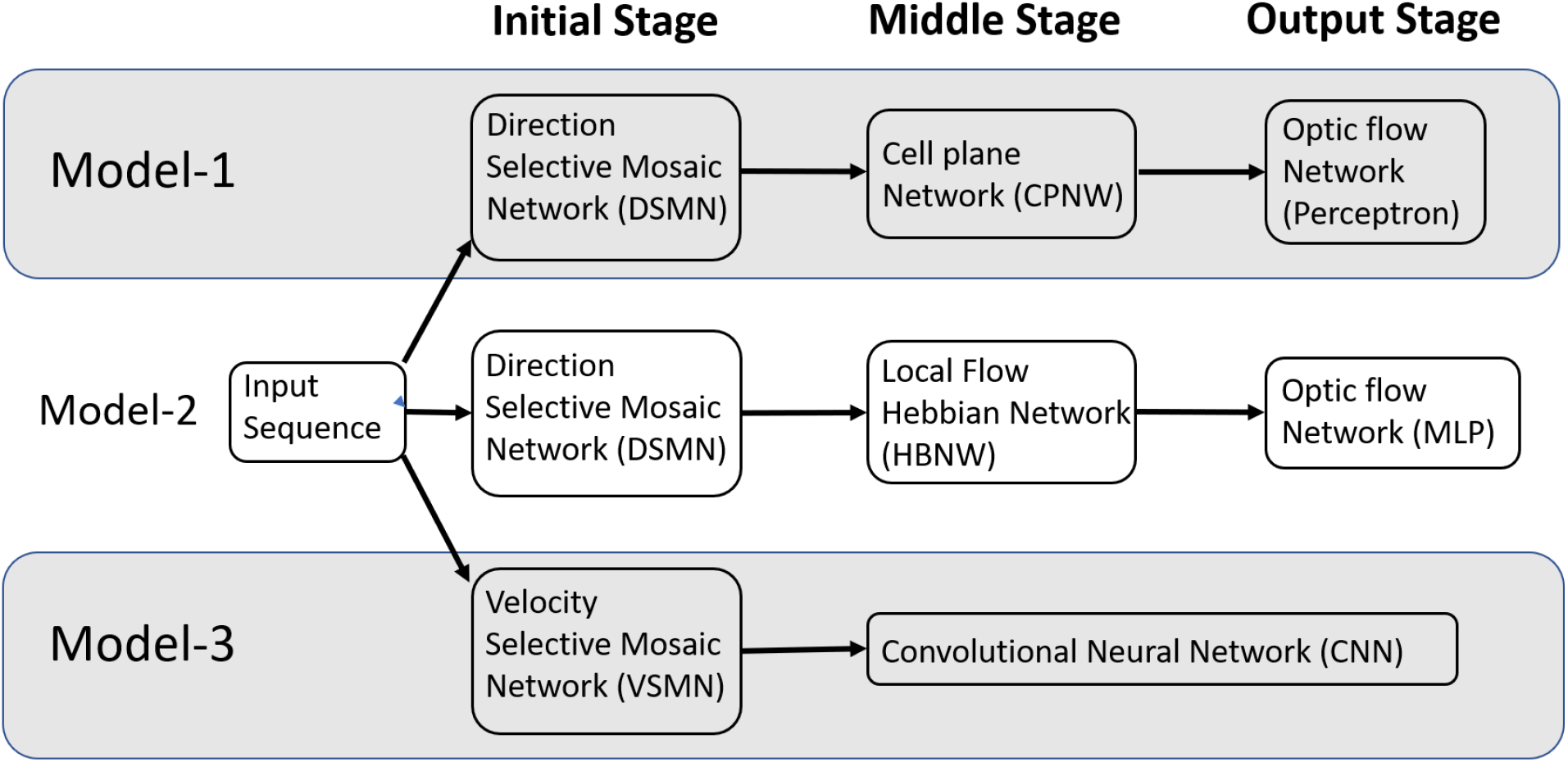
The multi-stage neural network models composed of the initial, middle and output stages that are simulating the neurons at V1, MT and MSTd respectively.

## 2. MODEL ARCHITECTURE AND LEARNING RULES

This section first describes various components/ sub networks used in all the three models and then presents the complete pipeline for all the three models. Before that we list out various physiological evidences used in designing these models.

### 2.1 Physiological evidences used in designing the model

Neurophysiological studies (Duffy & Wurtz, 1991a, 1991b; Graziano, 1990; Saito et al., 1986; Tanaka & Saito, 1989) have found that most of the neurons in the dorsal part of the medial superior temporal (MSTd) area of the visual cortex in the primates’ brain are responsive to different types of optic flow stimuli. It also has been found that MST receives strong projections from the middle temporal (MT) area (Maunsell & van Essen, 1983a, 1983b; Ungerleider & Desimone, 1986) where the neurons selectively respond to the orientation and velocity of the visual stimuli (Albright, 1984; Rodman & Albright, 1987). It is therefore natural to assume MT area to be the preprocessing stage to the optic flow processing taking place in MST area. There is physiological evidence that translational motion is computed in area MT (Movshon et al., 1985) while radial and rotational motions are first seen in the response properties of cells in MSTd (Duffy & Wurtz, 1991a, 1991b; Tanaka et al., 1989; Tanaka & Saito, 1989). Thus, according to this view, optic flow stimuli are processed serially, starting in the striate cortex with the analysis of motion in local parts of the visual field by direction selective cells with small receptive fields^1^. This local motion information is globally integrated in area MT by cells with larger receptive fields, which compute pattern motion - in this case translational motion. Finally, global radial and rotational motion is encoded by MSTd cells with much larger receptive fields based on their MT input. The selective responses of some MSTd cells are said to be position dependent, while those of others are position independent. In support to this view, it was suggested by several researchers (Saito et al., 1986; Tanaka & Saito, 1989) that the receptive field of an MST cell responsive to circular or radial motions is composed of a set of directionally selective MT cells arranged in accordance with the pattern of that optic flow component. Thus, the input MT cells would be arranged radially in the case of an expansion/contraction MST cell, or arranged circularly in the case of a rotation MST cell. Keeping these earlier proposals in view, to understand the responses of MST neurons and to explain how motion information is extracted to discriminate the type of the optic flow, we proposed an architecture composed of three stages: the initial stage consists of direction selective neurons that are trained to respond to the direction of motion of dots present in a given receptive field; the middle stage neurons are trained with translational sequences so that each neuron is selective to the direction of motion of local translational motion; neurons in the output stage are tuned to the type of the optic flow present in the input sequence. Compared to the algorithms of optic flow analysis proposed by computer vision community, the proposed modeling approach is more physiologically plausible and is able to account for some of the response properties of MSTd neurons. Fig. 1 shows the schematic representation of the three models proposed.

### 2.2 Direction selective mosaic network (DSMN)

The initial stage consists of a 16×16 mosaic of 2D arrays (“tiles”) of neurons, named Direction Selective Mosaic Network (DSMN). Every tile is an independent neural field (NF) wherein the neurons respond preferentially to the direction of motion of a dot present within their receptive fields. The neurons in a NF have lateral connections, thereby making the response of the NF neurons dependent on past history. The input to DSMN can be visualized as 16×16 non- overlapping image patches of size 5×5. Thus, the size of the input image is 80×80 (= (16×16) x (5×5)). Each NF is composed of 20×20 of neurons, receiving common input from a single 5×5 patch of the input image (Stage1: DSMN in Fig. 2). Although all the neurons in a given NF respond to the same 5×5 patch of their input image, they develop distinct selectivities, since they have random initial weights, and the initial differences are amplified by the competitive dynamics within the NF, as explained in more detail in the following section. As a result of training, NF neurons were clustered into various populations/ groups in such a way that each is selective to a specific direction of motion of a dot.

**Figure 2:**
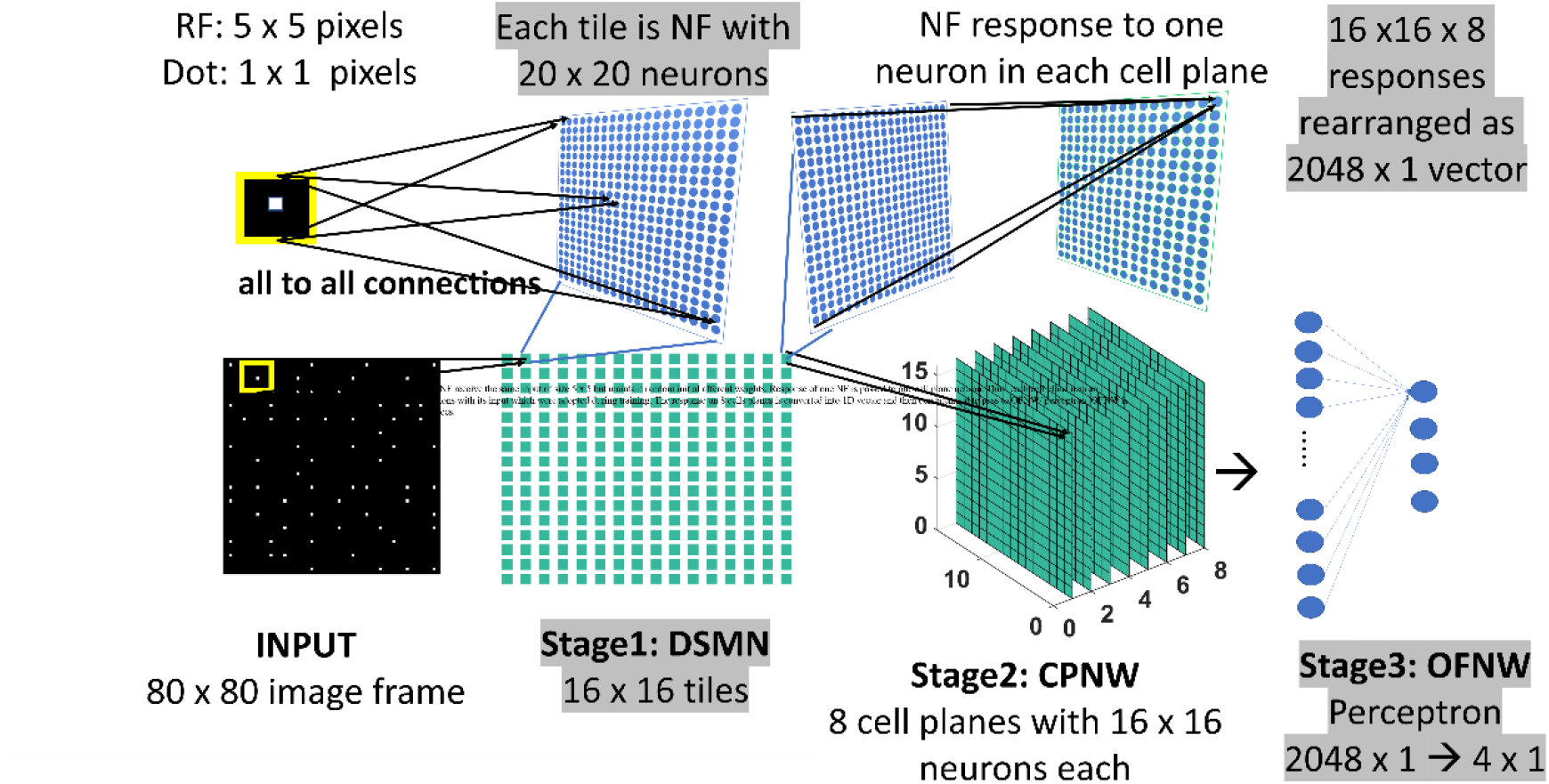
Model-1 architecture: All the neurons in NF receive the same input of size 5×5 but maintain random initial afferent weights. Response of one NF is passed to one cell-plane neuron. Thus, each cell-plane neuron maintains 400 (20×20) afferent connections with its input which were adopted during training. The response on 8 cells-planes is converted into 1D vector and then concatenated to pass to OFNW/ perceptron. OFNW is trained to classify 4 types of flow sequences.

#### Training procedure: the neural field (NF)

As shown above, the input image provided to DSMN is of size 80×80 pixels. Each NF is composed of 20×20 neurons. The neurons in a given NF receive common input from 5×5 window of input image. The initial response of NF neuron *(i, j)* is calculated as,

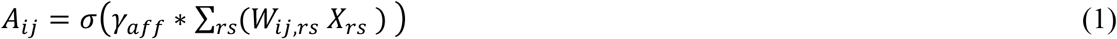

where *X* is 5×5 image window of neuron *(i, j). W* is afferent weight matrix of neuron *(i, j)*. Let *X*_*rs*_ be the pixel position in the image window then *W*_*ij,rs*_ is the afferent weight connection from *(r, s)* to *(i, j). γ*_*aff*_ is a constant scaling factor and is initialized before training begins. *σ* is piecewise linear sigmoid activation function.

The response of a NF neuron is influenced by both afferent inputs and the inputs from the lateral connections. Therefore, though the initial response is dominated by the afferent input, *A*_*ij*_, subsequently the response is further modified by the lateral connectivity of NF neurons. Lateral interactions are characterized by ON-center, OFF-surround neighbourhood. Two types of lateral connections exist: i) Excitatory laterals that connects neuron *(i, j)* with neuron *(k, l)* within a given neighbourhood. The excitatory neighbourhood is specified by the radius parameter *r*_*exc*_ and is initialized before training begins. The *r*_*exc*_ value is uniform for all the neurons within and across NFs. ii) Inhibitory laterals inhibit the response of the neuron *(i, j)*. It should be noted that the neuron *(i, j)* maintains inhibitory connections only with the neurons that are present outside the radius *r*_*exc*_ and inside the radius *r*_*inhb*_.

For several time steps *‘s’* (settling time), the response of the neuron *(i, j)* is modified by afferent and lateral interactions that take place simultaneously.

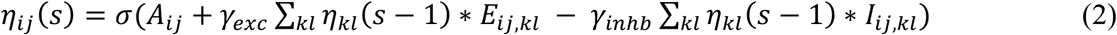

where *η*_*ij*_ stands for the activity of the neuron *(i, j), E*_*ij,kl*_ and *I*_*ij,kl*_ are excitatory and inhibitory weights from the neuron *(k, l)* to *(i, j)* that are randomly initialized before training begins. *A*_*ij*_, as defined in eqn. 1, is the total afferent input into neuron (*i, j*). The relative strengths of excitatory and inhibitory lateral effect are controlled by the constant scaling factors *γ*_*exc*_ and *γ*_*inhb*_.

At the end of ‘*s’* time steps, NF response settles down and all the three types of weights (afferent, excitatory laterals and inhibitory laterals) are updated. Let ‘*t*’ represent the time of presentation of current frame to NF with assumption that at *t=1*, frame-1 is presented. The afferent weight connections are adapted using symmetric Hebbian rule (eqn. 3) and the lateral weight connections are adapted using asymmetric Hebbian rule (eqn. 4).

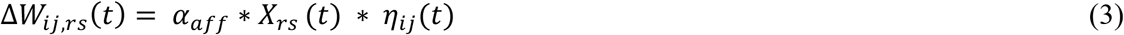

where *α*_*aff*_ is learning parameter for afferent weight connection. *X* is 5×5 image window from which neuron *(i, j)* receives input. *W*_*ij,rs*_ is afferent weight between the pixel position *(r, s)* and the neuron *(i, j). η*_*ij*_*(t)* is the activity of neuron *(i, j)* after the settling process for the current frame *‘t’*. The weights are updated after the presentation of each image in the input sequence as follows:

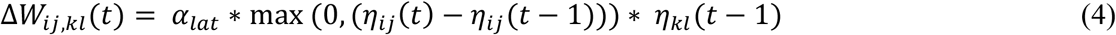

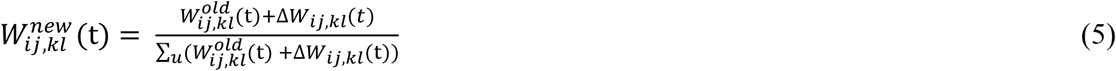

where *η*_*ij*_*(t)* is the settled activity on the neuron *(i, j)* produced in response to the current frame (the frame presented at time ‘*t*’), *η*_*ij*_*(t-1)* is the settled activity on the neuron *(i, j)* for the previous frame. *α* is the learning rate. Separate learning parameters (in place of *α*_*lat*_) were used for excitatory (*α*_*exc*_) and inhibitory (*α*_*inhb*_) connections. All the three types of weight connections are normalized separately as shown in eqn. 5 to prevent the neural activity from growing out of bounds. Various parameters used in the simulation were specified in Table1.

Note that the training procedure described above is for one tile (NF) that takes input from a 5×5 window of input image. 16×16 array of such tiles were trained sequentially one after the other by presenting an input image sequence consisting of 15 frames, each of size 80×80.

### 2.3 Cell plane network (CPNW)

The cell-plane, according to Fukushima (Tohyama & Fukushima, 2005), is a group of cells in which all cells have receptive fields of identical characteristics but the locations of the receptive fields differ from cell to cell. For example, a group of cells with same preferred moving direction is referred as cell-plane. The proposed CPNW consists of 8 cell-planes (Stage2: CPNW in Fig. 2), each responding preferentially to the translational motion of a particular direction. This translational motion selectivity of each cell-plane is achieved through training. For simplicity, and also for the fact that MT neurons are not sharply tuned for speed, only the direction of each flow field (DSMN response) was used to represent motion in MT stage.

#### Training Procedure

Each cell-plane consists of a 2D array of neurons of size 16×16; there are N_cp_ (=8) cell-planes used in model-1. Neuron *(p, q)* in the *n*^*th*^ cell-plane receives afferent input from all the neurons of *(p, q)*^th^ tile. Note that DSMN has 16×16 tiles, where each tile is made up of 20×20 neurons. The activity of the neuron *(p, q)* in the *n*^*th*^ cell-plane is computed using eqn. 6 and subsequently its afferent weights are updated using eqn. 7. All the initial afferent weights are set randomly. The eight cell-planes were trained sequentially one after the other by considering different stimuli set. In other words, cell-plane-1 is trained using stimuli set consisting of dots translated coherently in 0°, cell-plane-2 is trained using dots translated coherently in 45° and so on. Thus, each cell-plane is trained independently using dots moving coherently in 8 different directions. At the end of the training, the 8 cells-planes develop selectivities to 8 directions of dot motion.

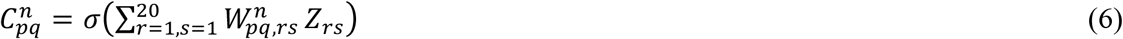

where *σ* is a sigmoid function. C^n^_pq_ is the response on the neuron *(p, q)* in the n^th^ cell plane. Z (20×20) is the *(p, q)*^th^ tile response in DSMN. W^n^_pq,rs_ represents the afferent connection from the neuron *(r, s)* within *(p, q)*^th^ tile to the *(p, q)*^th^ neuron in the n^th^ cell-plane.

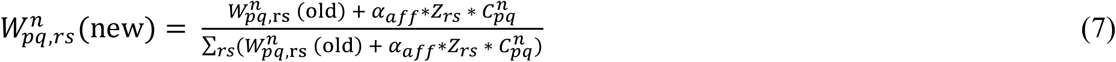

where *α*_*aff*_ is learning parameter, set to 0.05 during the simulation. Weights are updated for each sequence presentation. Updated weights are normalized to prevent them going out of bounds.

### 2.4 Hebbian network (HBNW)

The neurons in the Hebbian Network (HBNW) are arranged in similar fashion as CPNW neurons, but trained differently using competitive learning (Fukushima, 1988; Fukushima et al., 1997; Fukushima & Miyake, 1982). In this learning mode neurons compete in a winner-takes-all fashion, and the neuron receiving the largest input wins the competition and winner gets to modify its weights. Thus, neurons learn to respond to the inputs whose preferred direction best fits the local motion direction in the input. HBNW is composed of 16×16×8 neurons (Stage2: HBNW in Fig. 3) and is regarded as a 16×16 array of columns of neurons with 8 neurons in each column. A given column of neurons at location *(m, n)* receives a shared input Z_rs_ from a tile *(m, n)* in DSMN (that consists of 16×16 tiles). Similar to eqn. 6, the response of a neuron *(m, n, k)* is computed as a scalar product of the response of the tile *(m, n)* and the afferent weight matrix of the neuron *(m, n, k)* (eqn. shown again below the paragraph). All afferent weights are initialized randomly, accordingly the neurons across *(m, n)* column respond differently. The neuron in column *(m, n)* whose afferent weight matrix is closest to the input (which is the response of the tile *(m, n)* in DSMN) will produce highest activity, subsequently becomes a winner and its weights get updated following eqn. 8.

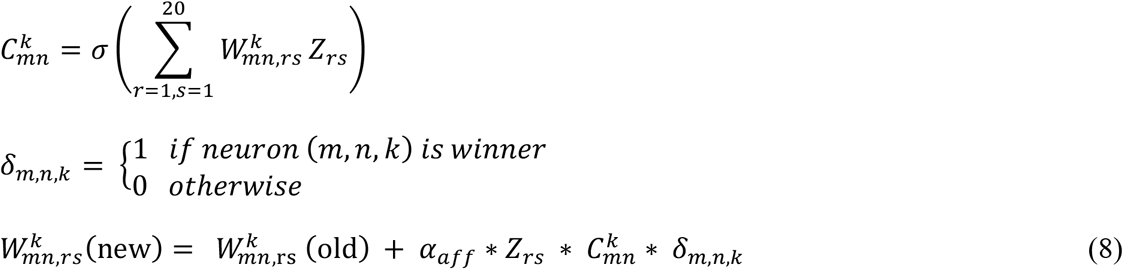

**Figure 3:**
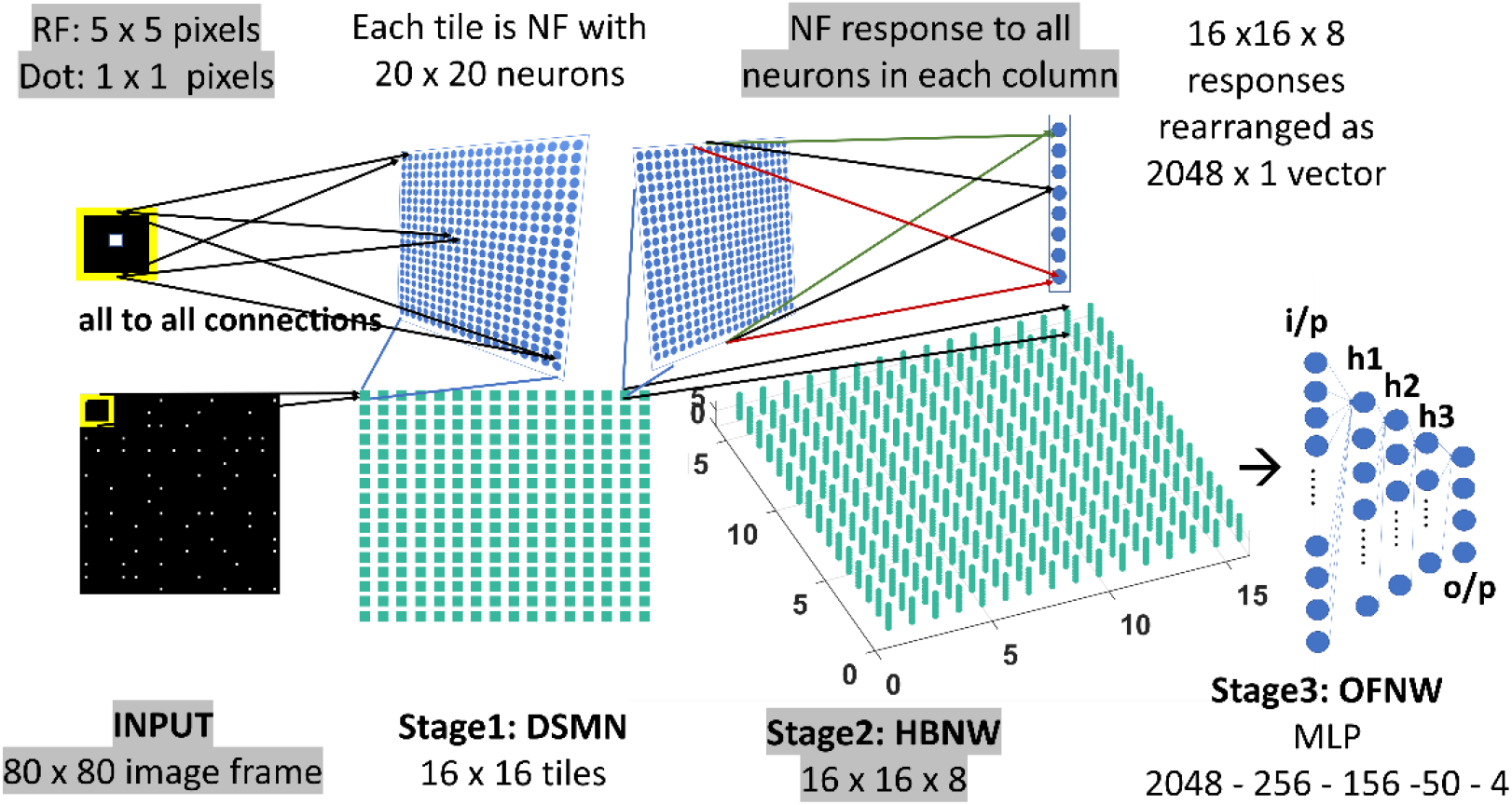
Model-2 Architecture: DSMN (consisting of 16×16 tiles) is similar to the one described in model-1. 16×16 NF responses were feed forwarded to 16×16 columns in HBNW. HBNW response is rearranged as 2048×1 vector before it is passed to the output stage-multi-layer perceptron (MLP) where the neurons are trained to recognize the type of optic flow present in the input sequence.

where *σ* is a sigmoid function. During weight adaptation, the afferent weights of the winner alone were normalized. C^k^_mn_ is the response of the neuron *(m, n, k)*. Z (20×20) is the *(m, n)*^th^ tile response in DSMN. Note that *(m, n)*^th^ tile is NF made up of 20×20 neurons. *Z*_*rs*_ refers to response on *(r, s)* neuron within NF in otherwards the tile *(m, n)* in DSMN. W^k^_mn,rs_ represents the afferent connectivity from the neuron *(r, s)* within *(m, n)*^th^ tile to the *(m, n, k)*^th^ neuron in HBNW. *α*_*aff*_ is learning parameter for afferent weights, whose value is set to 0.05 during the simulation.

#### Difference between HBNW and CPNW

The arrangement of the receptive fields of the neurons in both the networks are the same. CPNW neurons are grouped into 2D arrays and neurons in each group learns to respond preferentially to the same translational motion direction. Such grouping does not exist in HBNW, instead neurons in each column compete in a winner-take-all fashion. During training, different CPNW 2D arrays have access to different stimulus set. On the other hand, every neuron in HBNW has access to a continuum of input stimuli. Due to competitive learning, each neuron learns to discover a unique salient feature (direction of motion) present in the input. At the end of the training, a continuum of input stimuli is divided into a set of distinct clusters, where each cluster is represented by a particular set of HBNW neurons.

### 2.5 Optic flow network (OFNW)

A well-known perceptron and multi-layer perceptron with 3 hidden layers were used to implement the Optic Flow Neural Network (OFNW) in model-1 and model-2 respectively. Both the networks are developed and trained in MATLAB 2015.

#### Multi class perceptron

The multi class perceptron implemented as OFNW has an input layer followed by an output layer. The response of the CPNW is rearranged as a 1D vector before it is fed into the OFNW. The perceptron output layer consists of 4 nodes, each is trained to recognize the type of optic flow (expansion, contraction, clock wise rotation and anti-clock wise rotation) present in the given input sequence.

Let the training examples be *(X*_*1*_, *y*_*1*_*), (X*_*2*_, *y*_*2*_), …, *(X*_*n*_,*y*_*n*_), where *X*_*i*_ is an input vector and the labels *y*_*i*_ are drawn from the set *{l*_*1*_, *l*_*2*_,… *l*_*k*_*}*. Let the set of weight vectors to be learned are *{W*^*1*^, *W*^*2*^, …, *W*^*k*^*}*, then multiclass perceptron can be implemented as

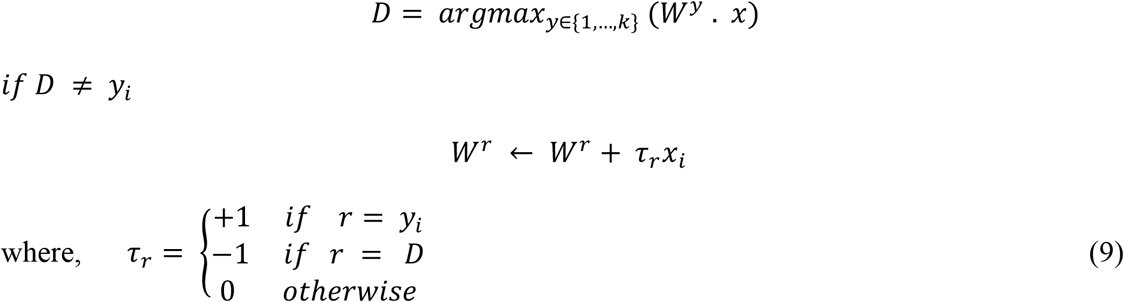

#### Multi-layer Perceptron

MLP network consists of 3 hidden layers, and is trained using regular backpropagation algorithm which was described in various studies (Bishop, 1995; Hornik et al., 1989; LeCun et al., 1992; Rumelhart & Hinton).

### 2.6 Velocity Selective Mosaic Network (VSMN)

Velocity Selective Mosaic Network (VSMN) used in model-3 (Fig. 4) is nearly the same as model-1 except it is formed out of 10×10 tiles and each tile is of size 48×48. Eqns. (1,3,4,5) are used for the calculation of VSMN initial response and weight adaptation. The equation for the calculation of settled response, eqn. 2, is modified. The input sequences generated to train the DSMN have fixed speed. Eqn. 10 is a modified form of eqn. 2 to make the network recognize variable speeds along with the direction of motion, i.e., to recognize velocity. In our simulations, we tried various scaling values for δ (see eqn. 12) ranging from 0.1 to 0.001. At higher values of δ the network fails to distinguish speed. At lower values of δ, the network response is unstable during the presentation of sequence. In other words, the lateral interactions could not produce unique activity patterns in NF to encode the speed feature.

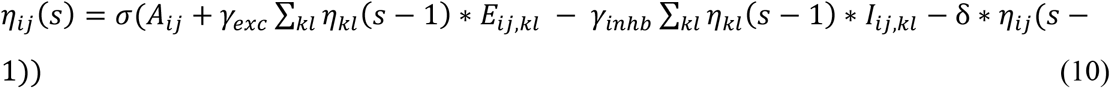

**Figure 4:**
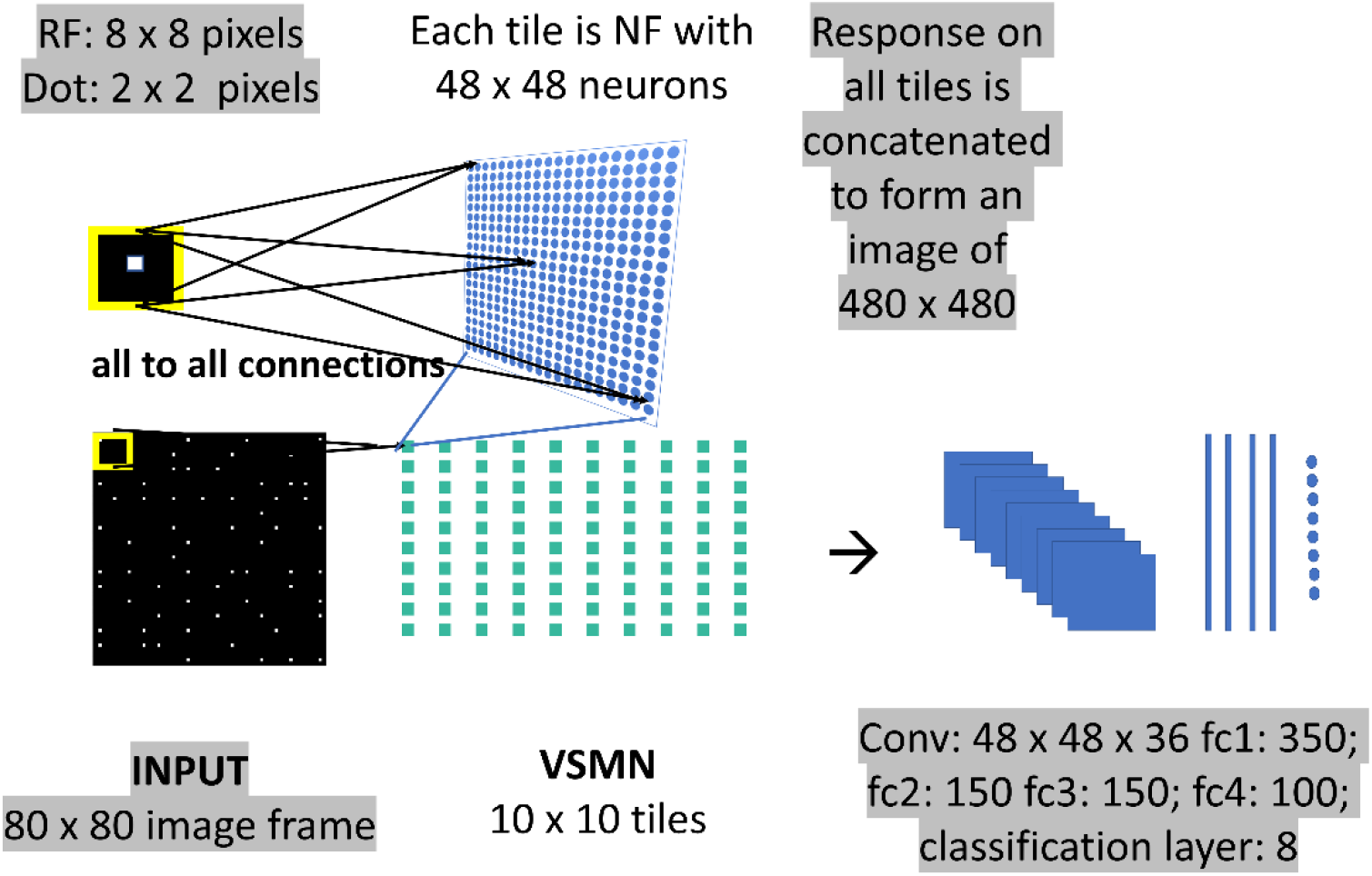
Model-3 architecture: The initial stage consists of VSMN that takes the input from image sequence. Each tile in VSMN is NF (48×48) where neurons are trained to encode direction and speed of a dot (2×2) within RF (8×8). The VSMN response obtained at the end of presentation of sequence is used to train CNN after concatenating the 10×10 tiles responses as a 2D image.

Unlike in DSMN where each tile is trained separately, in VSMN, each tile or NF is trained using input sequences (frame size 8×8) made up of 2×2 tiny squares moving in 8 directions and in each direction at 2 speeds. Thus, the training set of VSMN contains 16 sequences. The weights of a trained tile are copied to the remaining tiles in VSMN, to overcome the computational overhead.

### 2.7 Convolutional neural network (CNN)

VSMN response generated at the end of presentation of the entire sequence is used as input to the CNN. The responses on 10×10 tiles are concatenated to form an image of size 480×480. CNN is made up of one convolutional layer (with 36 feature maps) followed by 4 fully connected layers and a classification layer. CNN is trained to recognize type of optic flow along with its speed (8 classes = 4 flow types x 2 speeds). The design and simulation of deep network is carried out using MATLAB 2020a deep learning tool box.

### 2.8 Pipeline of all the three models

#### Model-1

As shown in Fig. 2, the three stages (DSMN, CPNW and perceptron) in model-1 are trained one after the other, in 3 steps, using the procedure described in the previous sections. In first step, each NF in DSMN is trained using the dot (1×1) moving in 8 directions such that the NF neurons develop direction selective responses. In second step, CPNW is trained with translational sequences moving in 8 directions (0, 45, 90, 135, 180, 225, 270 and 315°) such that neurons in each cell-plane responds maximally to specific translational motion direction. Note that while training CPNW, the DSMN weights are fixed and only its responses are forwarded. In the third step, the perceptron is trained to recognize type of the optic flow by using four types (zoom in, zoom out, clockwise, anti-clockwise) of optic flow sequences, while keeping DSMN and CPNW weights fixed.

#### Model-2

As shown in Fig. 3, model-2 consists of 3 stages-DSMN, HBNW and MLP and each is trained with moving dot sequences, translational sequences and optic flow sequences respectively in 3 steps similar to model-1.

#### Model-3

As shown in Fig. 4, model-3 consists of two subnetworks-VSMN and CNN. VSMN is trained using the dot (2×2) moving in 8 directions such that the NF neurons develop direction sensitive along with speed selective responses (i.e., velocity). These VSMN weights are fixed and only its responses are used while training the CNN. CNN is trained to recognize the type of flow (8 classes: 4 flow types x 2 speeds) present in the given input sequences. The training set of CNN is made up of dots that are allowed to make rotational (clock wise and anti-clockwise) and radial (inward and outward) trajectories.

For a clear understanding of all the 3 models the details about the preparation of training and test sets and the network performance were provided along with the results in Section 3.

### 2.9 Correlation measures used to construct response similarity matrix (RSM)

Pearson correlation measure (Emerson, 2015; Kpolovie, 2011) and Euclidean distance measures (Dokmanic et al., 2015) were used to construct response similarity matrices. The Pearson correlation coefficient *r*_*xy*_ is a statistical measure of the degree of linear correlation between the two variables *x* and *y. r*_*xy*_ takes values in the closed interval [−1, +1] (Emerson, 2015; Kpolovie, 2011). The value *r*_*xy*_ = +1 represents a perfect positive correlation between x and y, *r*_*xy*_ = −1 represents a perfect negative correlation between x and y, whereas the value *r*_*xy*_ = 0 indicates that no correlation.

### 2.10 Creating Input stimuli

#### Spatial Distribution

For model-1 and model-2, moving dot sequences were created by positioning 64 white dots on a black background of size 80 × 80 pixels with a density constraint that each 10 × 10 window typically accommodates only one dot. Each sequence is comprised of 15 frames. For model-3, dot stimuli were created by positioning 100 tiny white squares of size 2 × 2 pixels upon a black square grid of size 80 × 80 pixels with a constraint that each 8 × 8 window can accommodate only one tiny square at any given time. Each sequence is comprised of 10 frames.

#### Translational motion

Each dot configuration is moved (displacing x, y coordinates) in 8 directions (θ): 0°, 45°, 90°, 135°, 180°, 225°, 270° and 315°. The translational motion is incorporated in 15 or 10 frames as specified above, and each dot configuration adds 8 translational trajectories to the training set. If the dot exceeds the square boundary of the frame, it is wrapped around to reappear on the opposite side of the frame; thus, the dot density across the frames was kept constant. The horizontal and vertical displacement of a dot to incorporate translational motion is calculated by the eqn. 11.

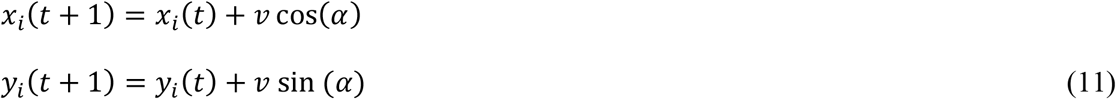

where α represents the direction of motion and the local speed is defined by ‘*v’*. In case of input stimuli for model-1 and 2, *v* takes only a single value (= 1) and for model-3 *v* takes two values (=1,2).

#### Optic flow motion

Each dot configuration is allowed to move along circular (clockwise, anti-clockwise) and radial (expansion, contraction) trajectories to create different flow sequences. Thus, each dot configuration adds 4 flow patterns to the training set. Let m and φ be the magnitude and orientation components of a dot at *(x, y)*. Then the trajectory of radial and circular motion is defined using the eqns. 12 and 13. Note that for radial trajectory magnitude (m) varies and for circular motion orientation (φ) varies.

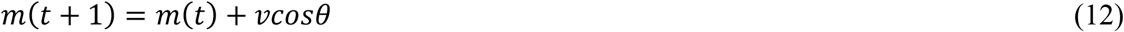

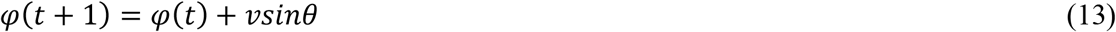

where *v* defines the local speed; *θ* defines the direction of flow, and takes the values 0 for expansion, π for contraction, -π/2 for clockwise rotation and π/2 for anti-clockwise rotation. Here also for model 1 and 2, *v* takes only a single value (= 1) and for model 3, *v* takes two values (=1,2).

## 3. RESULTS

### 3.1 Model-1

#### 3.1.1 DSMN response to translational dot sequences

Each tile in the DSMN is a NF. Each NF is trained with dot (1×1) moving in 8 directions (0, 45, 90, 135, 180, 225, 270 and 315°) from 3 different initial positions. Thus, training set is made up of 24 (8 directions x 3 positions) sequences. Trained NF response to the dot moving in eight directions and the corresponding direction selective map is shown in Figs. 5A, B. Even though all the tiles are trained to encode the direction of motion of a dot present within their receptive field, the neuron preferences across the tiles vary due to random initial afferent and lateral connections. That is, neurons at a specified location *(i, j)* in all the tiles, do not always respond to the same direction. Various network and learning parameters used in the simulation are given in Table 1. Fig. 5D represents the map of direction selectivity of DSMN. By comparing the color patches in the maps produced by each tile, one can understand that different neuronal populations are active in different tiles in response to the translational dot pattern moving in a specific direction. Each colored patch indicates the direction preference of the neuronal population. Fig. 5C represents the response of DSMN to the translational sequence moving in 8 different directions. The neural responses produced by different tiles in DSMN are concatenated and displayed in ‘Resp’ column in Fig. 5C.

**Table1:** Parameters used to train tiles (NF) in DSMN and VSMN

| Parameter | Tile in DSMN | Tile in VSMN |
| --- | --- | --- |
| NF Dimension | 20 x 20 | 48 x 48 |
| Receptive field | 5 x 5 | 8 x 8 |
| $r_{exc}$ | 2 | 2 |
| $r_{inhb}$ | 5 | 4 |
| $\gamma_{aff}$ | 1 | 1 |
| $\gamma_{exc}$ | 21.6 | 50 |
| $\gamma_{inhb}$ | 1 | 1.5 |
| $\alpha_{\text{aff}}$ | 0.05 | 0.05 |
| $\alpha_{\text{exc}}$ | 0.05 | 0.05 |
| $\alpha_{\text{inhb}}$ | 0.05 | 0.05 |
| Time step (s) | 10 | 10 |
| Epochs | 500 | 250 |
| Image size | 80 x 80 | 80 x 80 |

**Figure 5:**
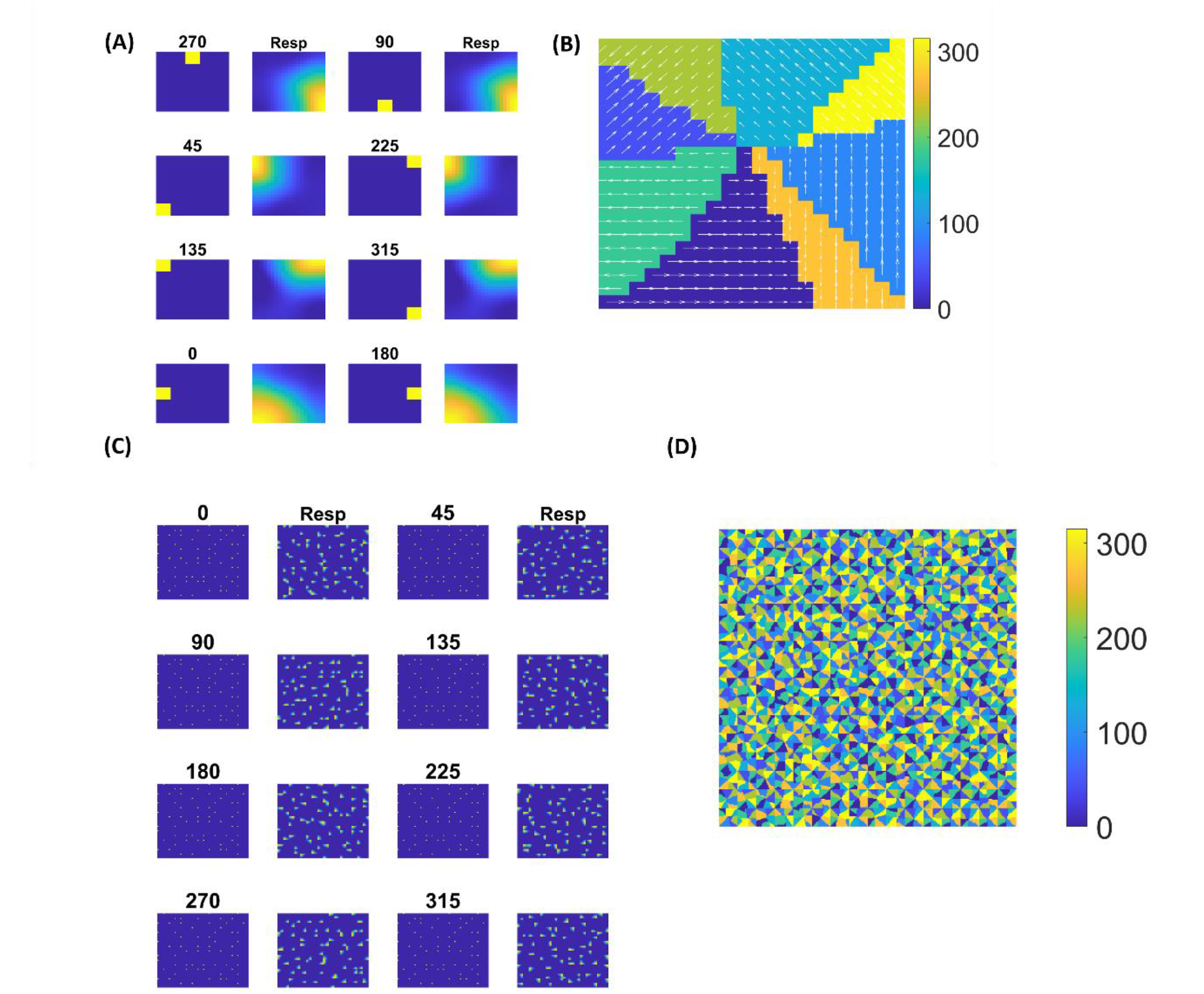
Neural field network and DSMN response: 1^st^, 3^rd^ columns in (A) display the first frame (5×5) of an input sequence and 2^nd^, 3^rd^ columns display the corresponding NF (20×20) activity. (B) shows the direction selective map for one tile. In (C) 1^st^, 3^rd^ columns show the frame (80×80) of a translational sequence and the response it elicits in DSMN (16×16 tiles) is plotted in 2^nd^ and 4^th^ columns. ‘Resp’ column shows the concatenated response in all the tiles of DSMN. (D) Displays DSMN’s direction selectivity map (concatenated direction selective maps of all 16×6 tiles). Each color on the map shows the neuron population with direction preference as specified in the color bar.

#### 3.1.2 Two-stage network response/ *CPNW response to translated motion*

Here we present a simulated results of the CPNW, which is the second stage of model-1. As described earlier, cell-plane network (CPNW) takes responses from DSMN, and is trained by repeatedly presenting translational motion sequences to DSMN (see section 2.3 and 2.8). Since the learning takes place in the different cell-planes independently (neurons in different cell-planes do not compete with each other) the responses of the cell-planes are similar but not identical. Here we used 8 cell-planes to encode 8 different motion directions. We could also choose to use more cell-planes to model a variety of MT and MST cells. 15 different initial dot configurations were translated in 8 directions to make 120 sequences, which were divided into training set (10×8 = 80) and test sets (5×8 =40).

The responses of the network stabilize after the training for 1000 epochs. Before the testing phase, the training set is presented to the CPNW and the winning cell-plane for each translational direction is recorded and is used as label to estimate CPNW performance on test set. Eight different cell-planes showed maximum responses to 8 different translational motion directions provided in the input stimuli. Now we present the test set consisting of eight translational motions of eight different directions (0, 45, 90, 135, 180, 225, 270 and 315°) that are not seen by the network before. As shown in Figs. 6A-H, each of the test sequence was responded to maximally and uniquely by one of the 8 cell-planes. We see that though each cell-plane responded most strongly to its preferred translational motion directions, it also responded to the neighboring directions with lesser intensity. CPNW showed 100% accuracy on test set.

**Figure 6:**
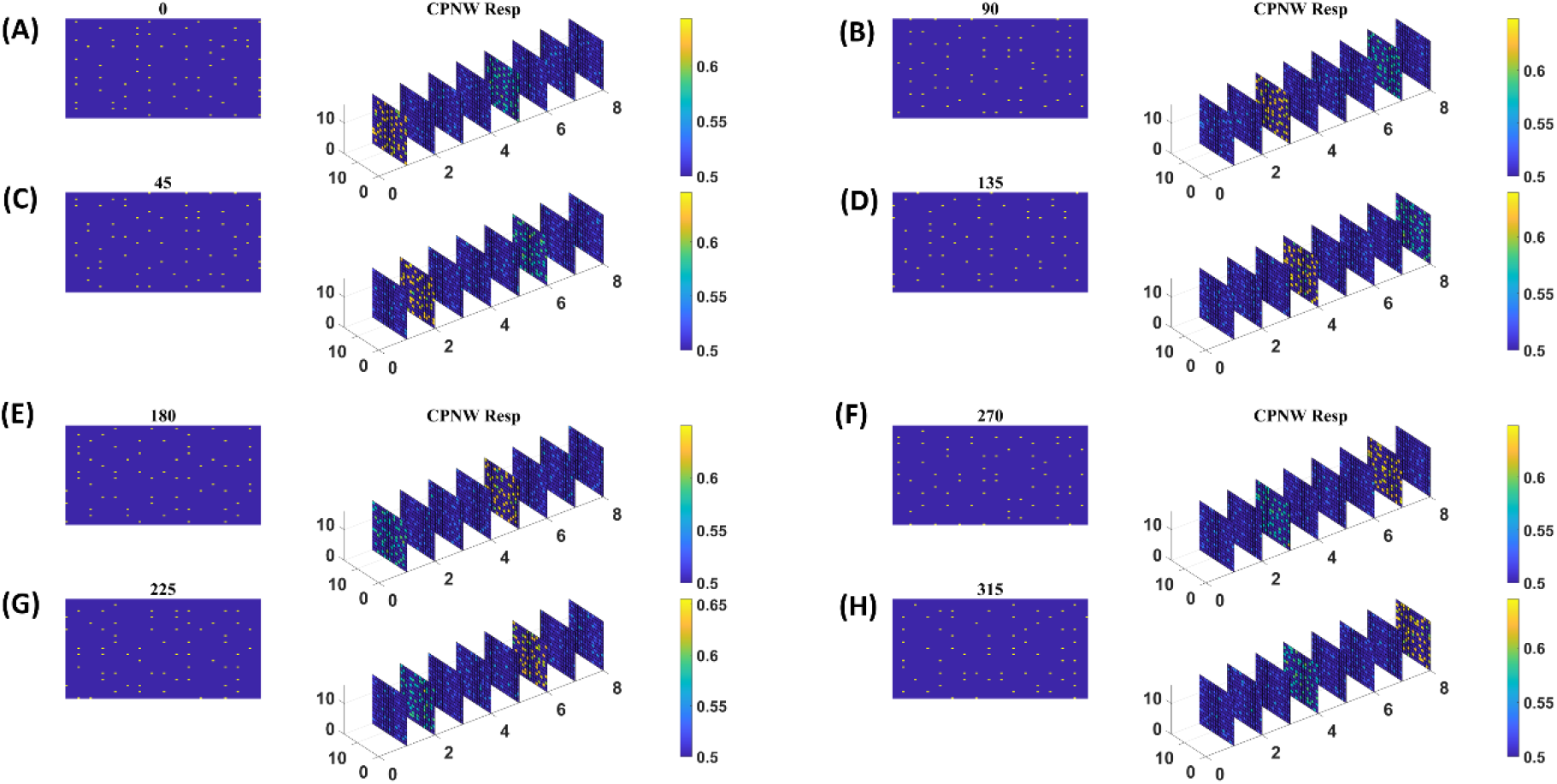
CPNW response to translational sequences: In all figures A-H a frame (80 × 80) of a translational sequence and its corresponding response on CPNW is displayed. The numbers 0, 45, 90 etc., represent the direction of motion of the input sequence. CPNW consists of 8 cell-planes, each shows maximum response to specific translational motion direction as result of training. The amount of activity produced by cell-plane neurons can be estimated using color bars. One can also observe that, though only one cell-plane produces highest activity in response to given motion direction (e.g., 0°), the cell-plane that encode opposite motion direction (180°) produce relatively high activity compared to the cell-planes that encode other motion directions.

#### 3.1.3 Three-stage network response/ perceptron response to optic flow sequence

Here we train the optic flow network (multi-class perceptron) simulating MST neurons and test the complete model-1 composed of all the three stages: initial DSMN, middle CPNW and output OFNW/ perceptron. The training set to train OFNW/ perceptron is composed of optic flow sequences including contraction, expansion, clockwise rotation and counter clock wise rotation each with 15 initial dot positions distributed over the image space. Thus, the training set is made up of 40 (10 × 4) flow sequences (see subsection optic flow motion in 2.10). We did not include planar translational patterns because here the main concern was the network’s response to different types of flow motion. The OFNW/ perceptron is trained by repeatedly presenting sequences in the training set in a random order to the DSMN. While training the perceptron, the weights of DSMN and CPNW that were trained earlier were kept constant and only their responses were feed forwarded to OFNW/perceptron. After 500 epochs, stable responses were obtained in OFNW/ perceptron. Now the test set comprising 20 sequences (5 initial dots x 4 flow types) is presented to the three-stage network (model-1) and the response is plotted in Fig. 7. Figs. 7A-D respectively shows the model-1 response to anti-clockwise, clockwise, expansion (Zoom Out) and contraction (Zoom In) motions.

**Figure 7:**
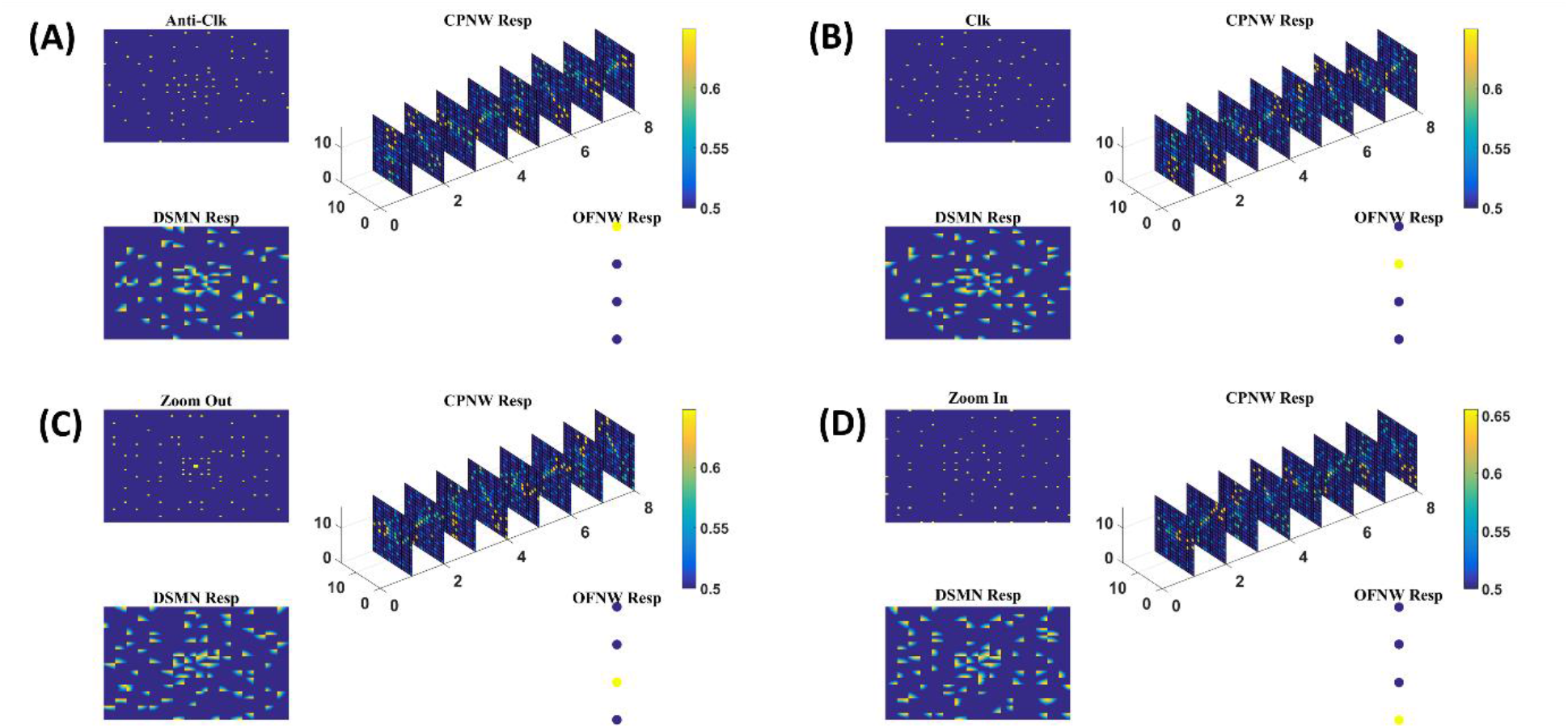
The Response of Model-1 to optic flow sequences: The response of OFNW to (A) Anti-clock wise sequence (B) clockwise sequence (C) zooms out or radially outward sequence (D) zoom in or radially inward sequence. In each case ‘DSMN Resp’ represents the populations of neurons that are active in each tile of DSMN. ‘CPNW Resp’ represents the subset of neurons that are active in each cell-plane in response to the given optic flow stimulus. The responses on 8 cell-planes are arranged as a 1D vector before giving it to OFNW. OFNW is a 4-class perceptron made up of 2 layers (input and output), and its response to 4 types of optic flow patterns is shown as OFNW Resp.

### 3.2 Model-2

#### 3.2.1 Two-stage network response/ HBNW response to translational motion

The first stage in model-2 is DSMN, which is the same as in model-1. The second stage, HBNW, is made up of 16 × 16 × 8 array of neurons. The training set used to train the CPNW in model-1, is now used to train the HBNW. As a result of training, the neurons in the HBNW learn to encode the local flow direction present in the small part of the image and on the whole continuum of neurons (16 × 16 × 8) encodes global motion information present in the input sequence. HBNW is trained by repeatedly presenting translational motion sequences to DSMN, whose responses in turn were forwarded to HBNW neurons. Training is carried out for 10000 epochs (learning rate = 0.05).

Trained HBNW responses to translational sequences of 180° are plotted in Fig. 8. Fig. 8A displays the first frame of translational motion sequence, Figs. 8B and C represent the corresponding DSMN and HBNW responses respectively. In Fig. 8C, one can observe that, at each (m, n) location along z direction, only one neuron shows the highest response (winner) ∼0.6 as indicated in the color bar; also, the winners at each vertical column are quite distinct. This is because a group of neurons along each vertical column that takes the same DSMN tile response, will be trained to recognize salient motion direction existing in the input pattern in the sense that similar set of input patterns (i.e., patterns having same translational motion direction) will always excite one particular neuron and inhibit all others. Thus, during the training process, due to initial random afferent connections one neuron from the group (m, n) shows the highest response, becomes winner and gets its weights updated. During the course of training, other neurons within the group will become winners when they encounter a different input. At the end of training, all the eight neurons within a group get tuned to 8 different directions. The arrows are plotted in Fig. 8C at each winner neuron to indicate their direction preferences. By the end of this competitive learning process, the continuum of input patterns is divided into a set of clusters, each cluster is represented by a particular population of the responding neurons. Note that each input pattern activates a large number of HBNW neurons, but the response selectivity is represented by the population of winner neurons. Thus, the different motion patterns presented to the two-stage network form a set of separable clusters in the feature space, which is typical for competitive learning. However, the class boundaries are not as crisp as in the case of CPNW.

**Figure 8:**
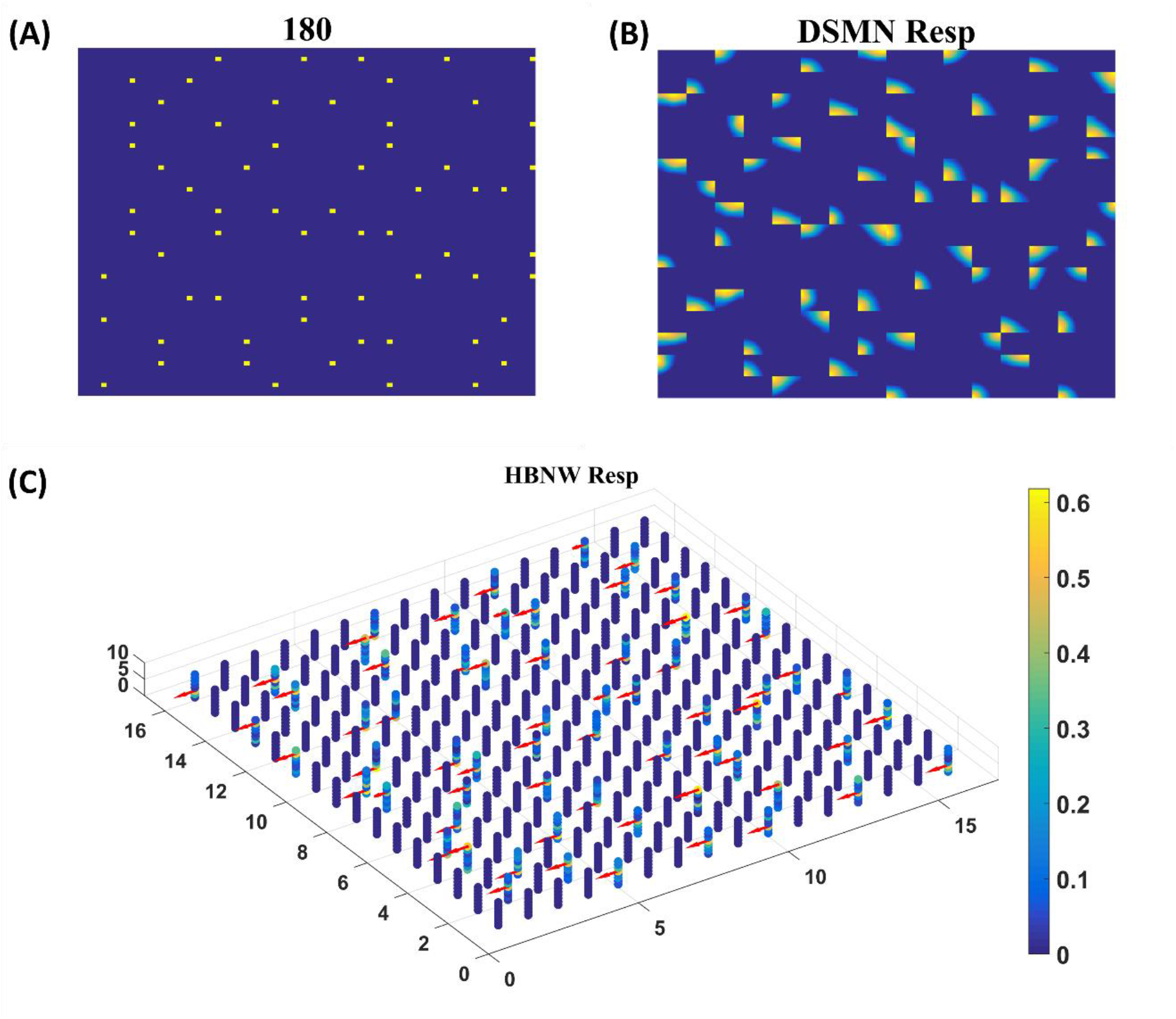
HBNW response to translational motion sequences: Here we plotted HBNW response to translational motion sequence: 180°. (A) represent a frame in an input sequence and corresponding DSMN response is shown in (B). Hebbian network (16 × 16 × 8) response titled as ‘HBNW Resp’ is plotted in (C). At each vertical column the winner is highlighted with the arrow whose head indicate the neuron’s direction preference.

Trained HBNW neurons show the highest response when their preferred direction best fits the local motion direction in the input. We observed the winner neuron responses by presenting all 120 translational motion stimuli. Eight winner neurons were observed in response to 8 motion directions along each vertical column. The direction preferences of HBNW neurons are plotted in Fig. 9. One can compare the winner neurons shown in Fig. 8C with the neuron preferences shown in Fig. 9. We observed that the HBNW neuron in each vertical column encoding eight different motion directions, without allowing emergence of redundant and dead neurons.

**Figure 9:**
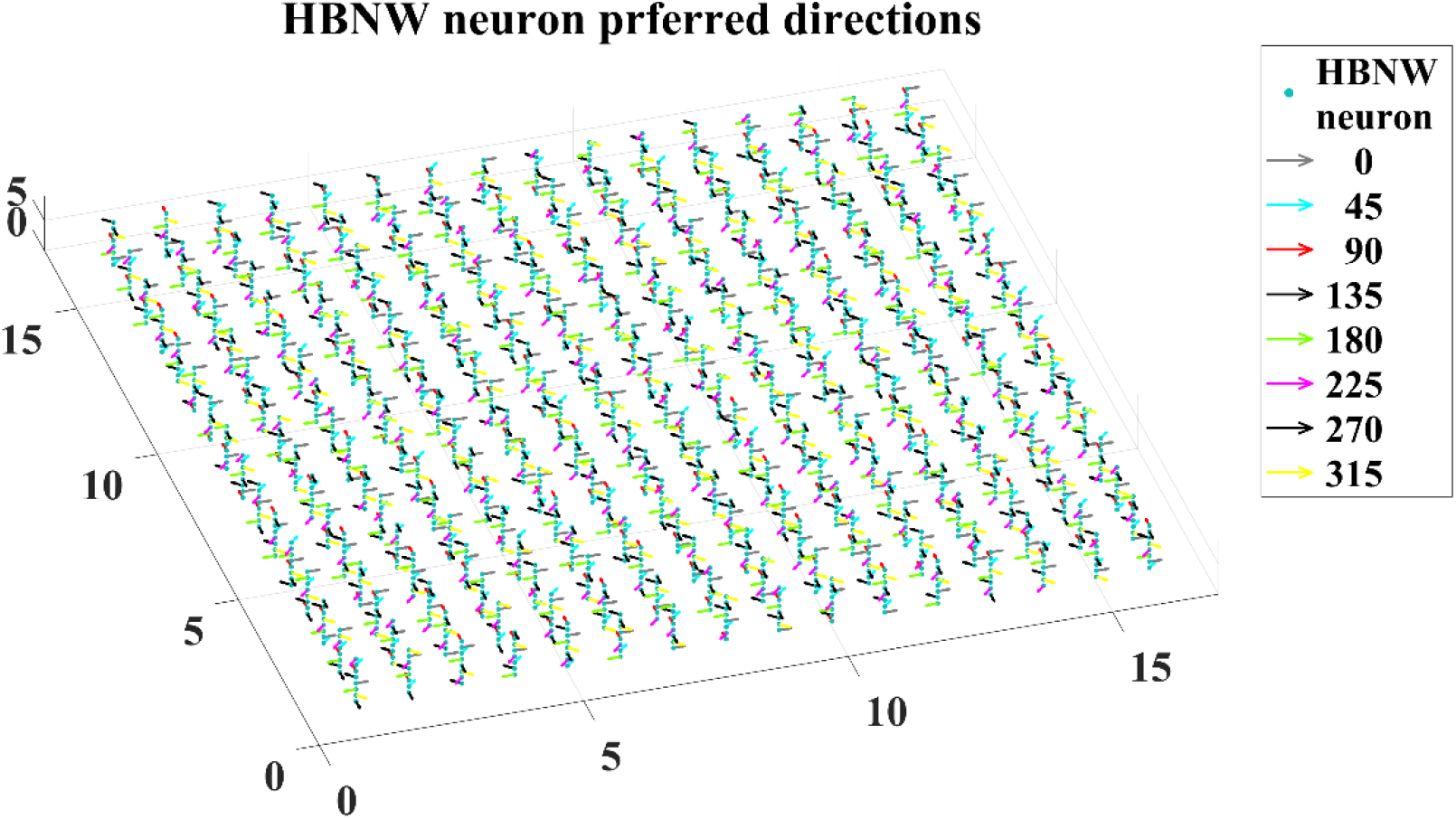
It shows the direction preferences developed by HBNW neurons in response to training (can be interpreted by observing coloured arrows more carefully).

#### 3.2.2 Three-stage network response/ MLP response to optic flow motion

Here we train the OFNW (multi-layer perceptron) simulating MST neurons and test model-2 composed of all the three stages: DSMN, HBNW and OFNW/ MLP. The MLP consists of input layer (2048 × 1), three hidden layers each of size 256 × 1, 156 × 1 and 50 × 1, and an output layer (4 × 1) (Fig. 3). It is trained using regular back propagation algorithm for 5000 epochs (learning rate = 0.1). The activation function used by nodes in the hidden and output layers are sigmoid and SoftMax respectively. Note that the neuron responses produced by CPNW is very different from the responses produced in HBNW during competitive learning.

As the nature of the input presented to output stage varies in model-1 and 2, different classification algorithms were proposed for OFNW. Similar to model-1 the training and test set for MLP in model-2 is composed of 40 and 20 flow sequences respectively each including contraction, expansion, clockwise rotation and anti-clockwise rotation. While training MLP, DSMN and HBNW weights were kept constant. The accuracy obtained on training set and test set are 100% and 95% respectively. MLP recognized one anti-clockwise motion sequence in test set as clockwise one. Once MLP training is completed, the response of the three-stage network is observed for every sequence in test set. The response for one ‘Zoom In’ sequence is plotted in Fig. 10 where ‘OFNW resp’ represents the activity of the output layer neurons in MLP.

**Figure 10:**
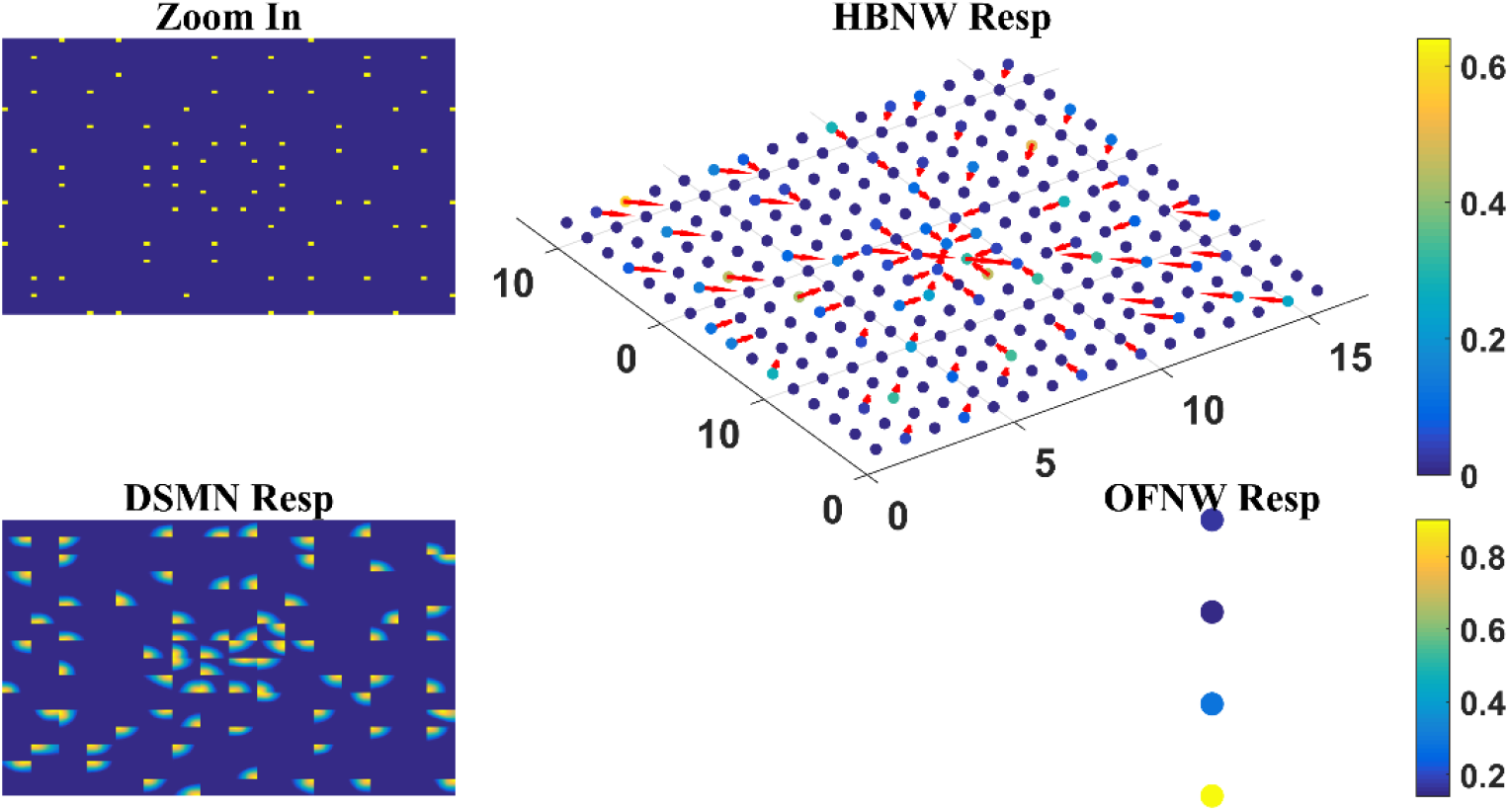
Model-2 responses to a ‘Zoom In’ optic flow sequence: ‘Zoom In’ represents frame in a radially inward flow. ‘DSMN Resp’ represent the populations of neurons that are active in each tile to the input sequence. Hebbian network response (top view) is plotted under ‘HBNW Resp’. Each column is represented by a winner along with its direction preference (arrow). ‘OFNW Resp’ represents the response of the classification layer nodes in MLP, each node is encoding specific flow type.

### 3.3 Model-3

The model-2 is biologically plausible optic flow recognition model, in which training of initial, middle and output stages is designed based on the knowledge of the response properties of the neurons present at various levels of motion pathway. On the other hand, in recent years, data driven CNNs, developed based on early studies of the visual system (Schmidhuber, 2015), have been widely used as visual encoding models. These encoding models work by establishing a nonlinear mapping from visual stimuli to features, and a suitable feature transformation is critical for the encoding performance (Tang et al., 2018). Studies have also shown that a deep network is comparable to the human visual system, which can automatically learn salient features from large data for specific tasks (Agrawal et al., 2014; Cohen et al., 2017). Recent studies by Güçlü and van Gerven demonstrated the similarity between CNN and visual pathways, revealing a complexity gradient from lower layers to the higher layers (Güçlü & van Gerven, 2015, 2017). On similar lines we developed data driven optic flow recognition model (presented in Fig. 1D) using CNNs.

#### 3.3.1 VSMN tile response

In this study, we investigate whether CNN can serve as a model of the macaque motion-processing network. There is evidence that, unlike V1 neurons, a subset of neurons within primate extrastriate cortex (MT) appear broadly tuned to local image speed and direction (Maunsell & Newsome, 1987; Movshon et al., 1985). The model is said to be biologically plausible if and only if V1, MT and MST representative neurons in the model show response properties analogous to those of real V1, MT and MST neurons. Currently the middle stage (CPNW, HBNW) neurons in model-1 and model-2 estimate local flow motion purely based on the direction selective responses from DSMN. To simulate MT-like responses to the local image motion according to the product of their direction and speed responses, we created Velocity Selective Mosaic Network (VSMN) in place of DSMN, which encodes speed together with direction.

Model-3 consists of VSMN followed by CNN (Fig. 4). Each tile in VSMN is a NF and is trained to recognize the velocity of the input stimuli as described in section 2.6. Fig. 11A shows the response of a tile in VSMN for the input stimuli consisting of 2 × 2 dot moving in 8 directions. Here dot is allowed to move one pixel and two pixels ahead for each time step that constitute two speeds. Thus, input stimuli consisting of 16 inputs (8 directions and 2 speeds). Fig. 11B plots the velocity selective map consisting of 16 populations. Each population is highlighted with a colored arrow (red for speed1 and black for speed2), indicating the preferred speed and direction of the motion.

**Figure 11:**
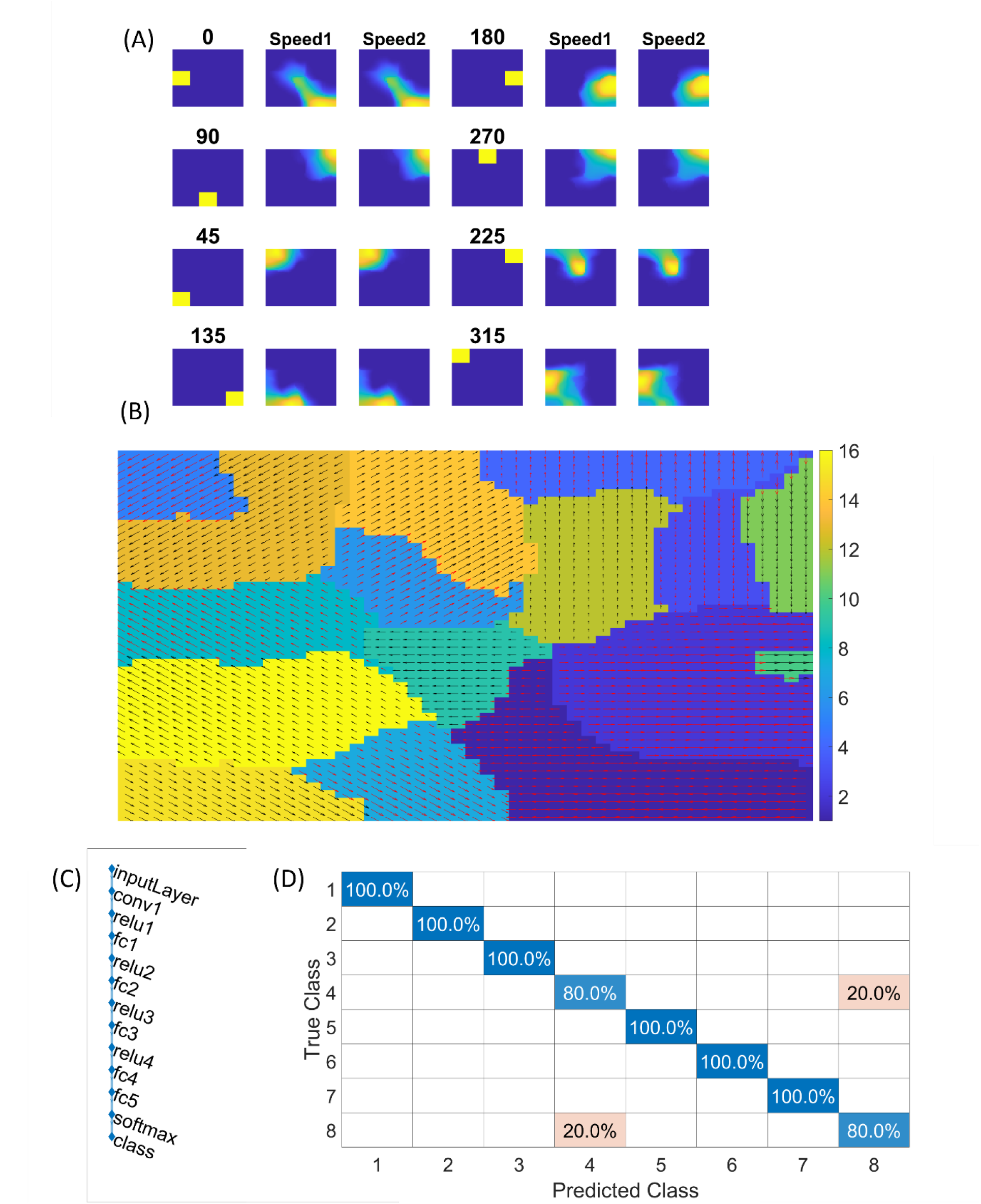
NF response in VSMN: In (A) 1^st^ and 4^th^ columns display the first frame (8×8) of an input sequence that consists of dots (2×2) moving in specific direction with two speeds. The 2^nd^, 3^rd^, 5^th^ and 6^th^ columns represent the corresponding NF (48×48) activity. The same neuron population become selective to the inputs of same direction and different speeds, however within the population neurons have different speed preferences which are shown in (B)-the velocity selective map. Neuron direction preferences are shown by head of the arrow and the speed preferences are shown by the arrow color. Color bar indicate 16 input types (8 directions x 2 speeds). 1 to 8 belongs to speed-1 and 9-16 to speed-2. (C) shows the architecture of CNN. (D) shows the confusion matrix obtained for the test set.

#### 3.3.2 Training CNN with optic flow sequences

CNN is trained with 8 classes. Thus, the training stimuli for CNN is made up of four types of flow sequences (expansion, contraction, clockwise rotation and anti-clockwise rotation) and each occurs at 2 speeds. A total of 80 motion sequences (10 initial dot positions x 4 flow types x 2 speeds) were generated for the training set and 40 (5 × 4 × 2) sequences for the test set. The CNN is trained to recognize the type of optic flow present in a given input sequence, by repeatedly and randomly presenting input from the training set to its previous VSMN, whose final response is forwarded to the CNN. The architecture of the CNN and various learning parameters used to train the CNN are described in Table 2. Note that the CNN is trained only on optic flow stimuli and not exposed to the translational stimuli. Fig. 11C and D indicate the CNN architecture, the confusion matrix and the learning curve respectively. For the implementation of CNN and to visualize the CNN feature maps, we used MATLAB with Deep Learning Toolbox. The trained CNN gave classification accuracy-100% for training data and 90% for held-out test data.

**Table 2:** CNN architecture and learning parameters used

| Layer | Size |
| --- | --- |
| Convolution Layer (conv1) | 48 x 48 x 36 |
| Fully connected layer (fc1) | 350 x 1 |
| Fully connected layer (fc2) | 150 x 1 |
| Fully connected layer (fc3) | 150 x 1 |
| Fully connected layer (fc4) | 100 x 1 |
| Output layer | 8 x 1 |
| **Learning Parameters** |  |
| Learning rate | 0.01 |
| Batch size | 20 |
| Epochs | 350 |

#### 3.3.3 CNN response to Translational and flow sequences

Next the trained CNN is presented with both translational and optic flow sequences, and the responses are plotted in Fig. 12. ‘Conv1’ layer consists of 36 feature maps (Fig. 4). All these feature maps are arranged in a grid-like structure and plotted as shown in ‘Conv1 Resp’. Figs. 12A, B plots the conv1 response to 4 types of optic flow stimuli moving with speed-1 and speed-2 respectively. Whereas Figs. 12C, D and Figs. 12E, F display the conv1 response to translational stimuli moving 8 directions with speed-1 and speed-2 respectively. Note that the CNN network was never exposed to translational patterns during training. Still diffuse and sparse response patterns could be observed under conv1 response column, in response to translational and flow sequences respectively. One can compare these responses with the responses of CPNW in Fig. 6 and Fig. 7, where we obtained similar diffuse responses to translational sequences and sparse responses to optic flow sequences.

**Figure 12:**
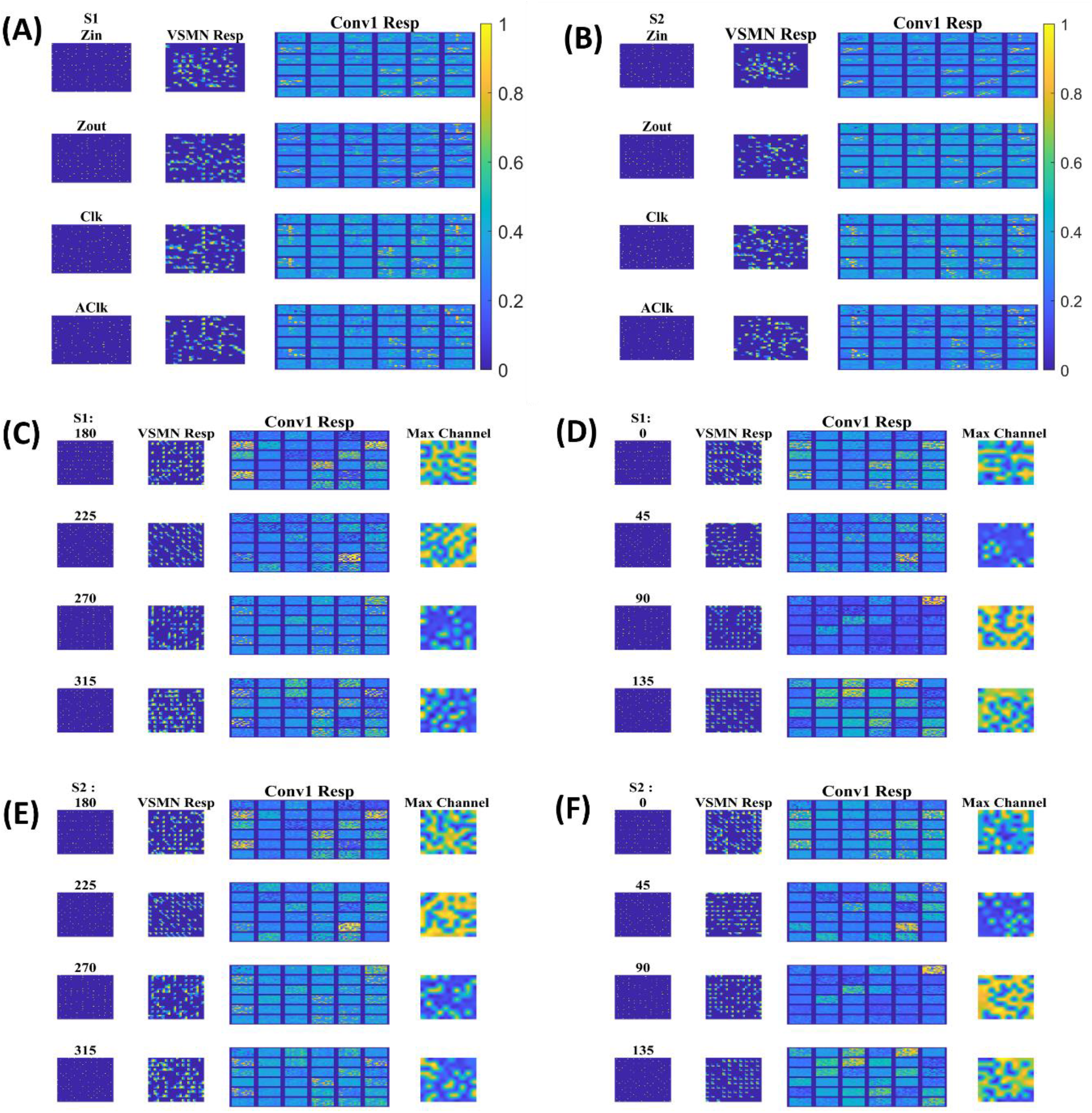
Model-3 responses to optic flow and translational motion sequences: In (A), (B) 1^st^ and 2^nd^ columns display the frame of an optic flow sequence and the corresponding VSMN response respectively. ‘Conv1 Resp’-indicates the activity on 36 feature maps of convolution layer. In (C), (D), (E) and (F) 1^st^, 2^nd^ and 3^rd^ columns display the frame of a translational sequence, corresponding VSMN resp and the convolution layer activity respectively. Out of 36 feature maps, the channel with the maximum response is plotted in the 4^th^ column as ‘Max Channel’.

#### 3.3.4 Development of translational motion selectivity and speed selectivity in conv1

We may now ask whether the CNN trained on optic flow motion sequences developed selectivity to translational sequences in its lower conv1 layer. If that is the case, conv1 layer is analogous to MT and CNN’s output layer is analogous to MST. To verify the above hypothesis, we presented the trained network with a test set, consisting of 240 translational sequences moving in 8 directions, with 2 speeds and started from 15 different initial dot configurations (15 × 8 × 2 = 16 classes with 15 sequences in each class). For all 240 inputs, the conv1 feature map or a channel with a maximum response is noted and the bar graph against each class is displayed as shown in Fig. 13A.

**Figure 13:**
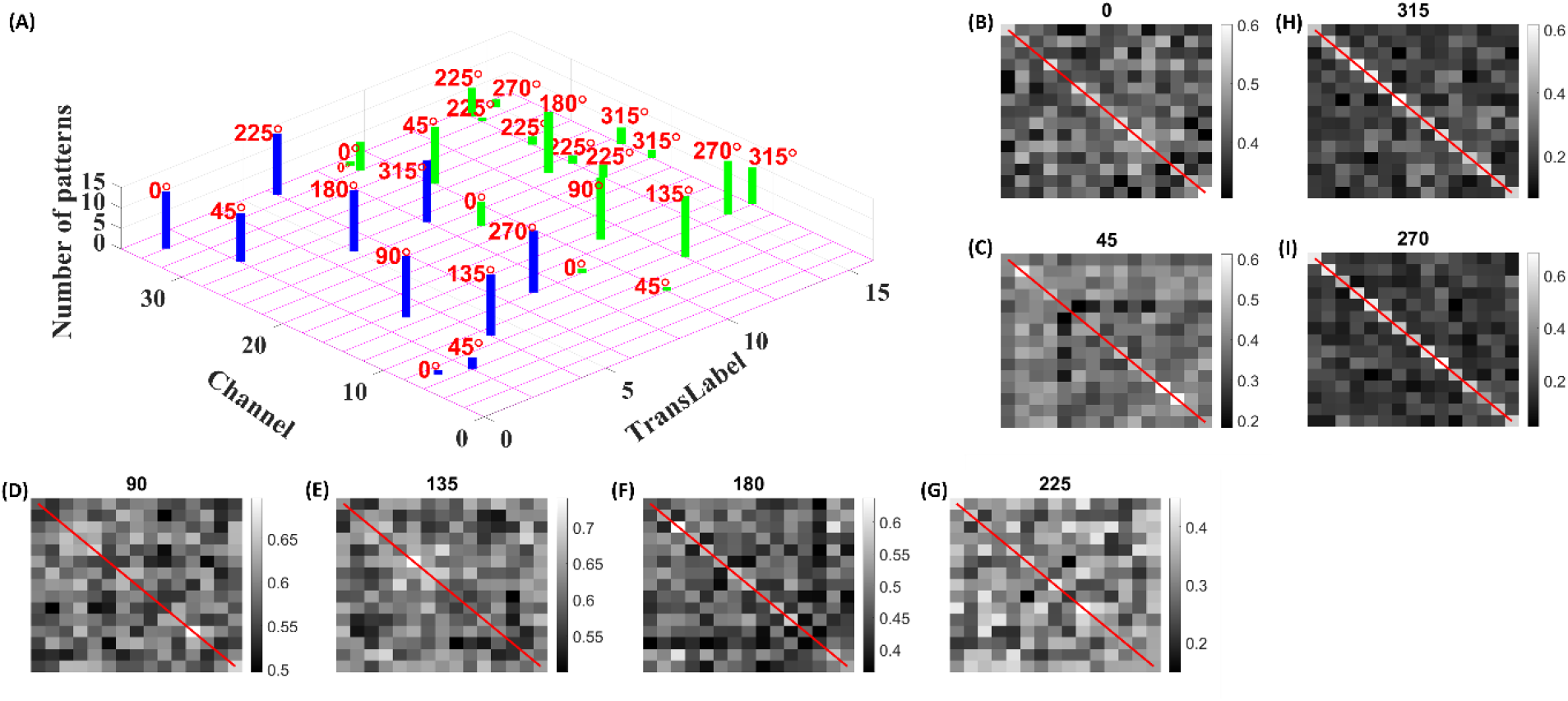
Development of speed selectivity in Conv1 layer: Each bar in (A) indicates the active channel corresponding to each input class (16 classes= 8 directions x 2 speeds; test set consists of 240 sequences: 15 initial dot positions x 16 classes; labels 1-8 belong to speed-1 inputs and are indicated by blue bars; labels 9-16 belong to speed-2 inputs and are indicated by green bars). For example, 0° translational sequence with speed-1 activates two channels 7 and 34. However more inputs activate channel 34 as indicated by bar length. One can observe that different channels encode different motion directions. However, inputs moving in same direction with different speeds seems to activate the same channel in bar graph. **Speed selectivity in Conv1 layer using RSMs**: (B) – (I) represent the response similarity matrices (RSMs). The matrix entries (15×15) indicate the pairwise correlation coefficients for speed-1 and speed-2 translational motion sequences whose direction is indicated on top of each figure. The diagonal elements of a matrix (as highlighted with red line) represent correlation coefficient calculated for the pair of sequences having same initial dot configuration and moving with different speeds. Most of the RSMs have no strong block diagonal structure (except for 315° and 270°), indicating that different neuronal populations in conv1 respond to different speeds. For 315° and 270° block diagonal structure can be seen due to the very low correlation values, not because of similar responses of conv1 channel to both speeds, as the high correlation value is 0.6 as indicated in color bar. Thus, even though the same channel codes for a sequence with different speeds, within the channel different subpopulations code for different speeds.

As shown in Fig. 13A different channels respond maximally to different translational directions, and also more than one channel showed highest activity to the same motion direction. These results are consistent with previous reports of a high degree of direction selectivity in MT with nearby units having similar preferred directions (J. H. Maunsell & van Essen, 1983a). These studies also reported that the MT units were sharply tuned for the speed of stimulus motion. However, the same channel in conv1 layer seems to become highly active to the inputs with same translational direction but moving at different speeds (as indicated by bars: blue-speed-1 and green-speed-2). MT neurons must have different speed characteristics to be consistent with the MT studies (Lagae et al., 1993). Orban et al (Orban et al., 1981) grouped cells in areas 17 and 18 into four distinct classes based on the broadness of speed tuning curve and the speeds to which they responded. To understand whether the conv1 layer developed speed selectivity as a result of training, we computed correlation matrices and plotted as shown in Figs. 13B-I. Initially, conv1 responses for all 240 translational inputs are obtained. For each direction, Pearson correlation between each pair of speed-1 and speed-2 (15×15 pairs) is calculated. The diagonal elements of a matrix (as highlighted with red line) represent the correlation coefficient measured between the translational patterns moving at two different speeds (speed-1 and speed-2) starting with same initial dot configuration. Smaller correlation values, as shown by the color bar and lack of block diagonal structure in plots (B-I), indicate that conv1 neurons display different responses to different speed stimuli, which is consistent with the cell properties in MT (Duffy, 1998; Duffy & Wurtz, 1991a, 1991b, 1995; Graziano et al., 1994; Graziano, 1990). Further to quantify this speed selectivity by conv1 channel, we trained a two-class linear classifier for each case displayed in plots (B-I). The accuracies obtained on test set for each direction 0, 45, 90, 135, 180, 225, 270 and 315° are 80%, 70%, 90%, 80%, 80%, 70%, 90% and 100% respectively. Thus, when CNN is trained using optic flow motion stimuli, the lower layers develop selectivity to translational motion which is analogous to MT.

We also calculated the Euclidean distance between conv1 responses to each pair of 8 translational directions, and for each speed separately. As shown in Fig. 14 (green lines), the responses to dot patterns moving in opposite directions (0°-180°, 45°-225°, 135°-315°, 90°-270°) have a higher correlation compared to the other pairs. Similar responses were seen while comparing the responses to speed-1 and speed-2 stimuli set. Thus, translational patterns with opposite motion directions develop similar activity pattern on conv1 layer, suggesting that they are closer in feature space.

**Figure 14:**
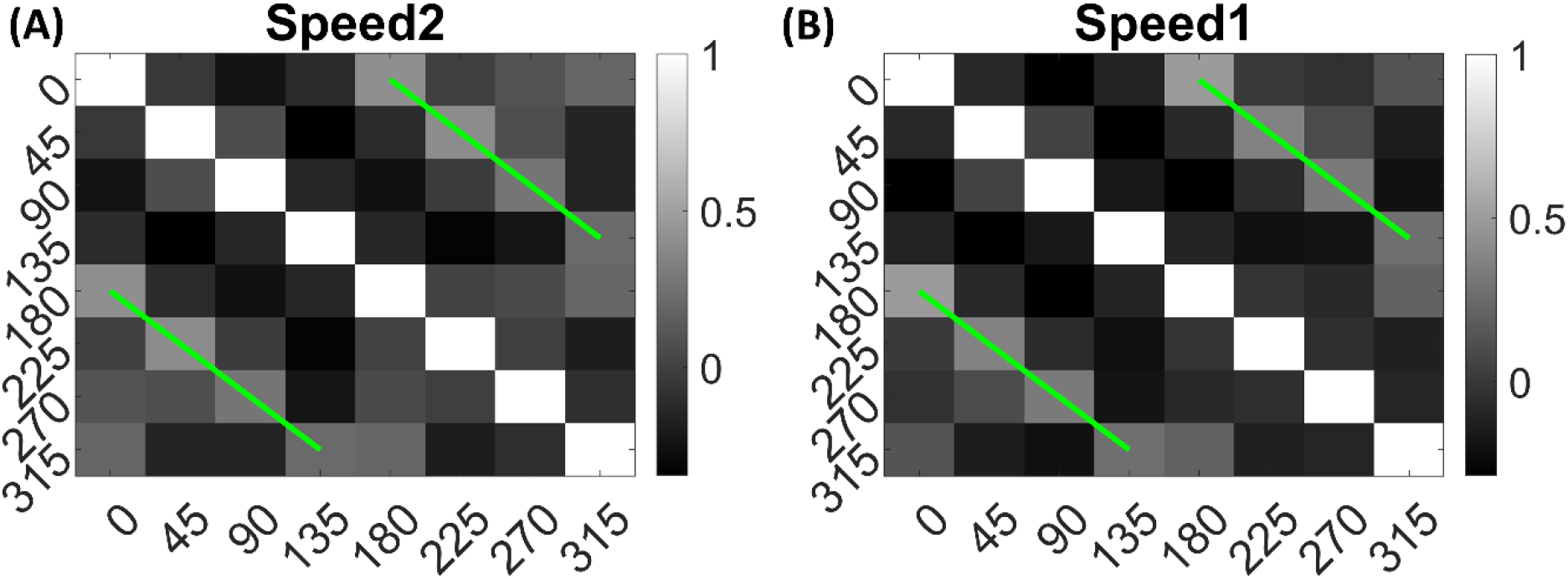
Direction tuning in conv1 layer: Each plot shows conv1 response similarity matrix (8×8) for each speed. Entries in the matrix indicating pairwise Euclidean distance between responses to translational motion directions (one sequence considered for each direction). The directions are indicated along the axes. From the plot one can observe that the conv1 layer produce similar response to the sequences with opposite directions. Matrix entries along green lines are positively correlated and the entries between orthogonal angle are negatively correlated (values are < 0).

#### 3.3.5 Correlating CNN responses with known properties of motion processing hierarchy

To investigate whether the CNN can explain known tuning properties of the macaque motion processing network (Duffy & Wurtz, 1991a, 1991b, 1995; Graziano et al., 1994; Graziano, 1990; Maunsell & van Essen, 1983a, 1983b), we calculated the responses of CNN layers (conv1and fc4) to translational (240) and flow (120) sequences made out of training together with test sets. We then calculated the correlation between the population responses to each pair of these sequences and constructed a population Response Similarity Matrix (RSM) of such correlations for all pairs.

Fig. 15 shows RSMs for convolution layer (conv1) and last hidden layer (fc4), for both translational motion and optic flow motion. In Figs. 15A, B, C and D the translational sequence numbers are grouped according to the motion direction (each forming sub-matrices of size 15×15). In Figs. 15 E, F, G and H the optic flow motion class numbers represent the type of the flow (here also each sub-matrix is of size 15×15). Each of 15 × 15 response similarity matrices indicate the pairwise correlation coefficients calculated for sequences made out of 15 initial dot configurations. In conv1 layer, the RSMs corresponding to the translational stimuli (Figs. 15A and B) show that different conv1 neuronal populations show selectivity to a different translational motion direction. In Fig. 15A and B the strong block-diagonal patterns can be seen, indicating that the populations show selectivity to specific direction of motion irrespective of initial dot position. Also, RSM entries between opposite motion directions are positively correlated, whereas RSM entries between orthogonal pairs are negatively correlated, indicating that opposite motion directions at the input space are arranged more closely at the feature space than the orthogonal motion directions. Figs. 15C and D display the RSMs corresponding to the translational stimuli in layer fc4 (last hidden layer). Even though a weak block-diagonal pattern can be seen, the clear distinct response profiles to various translational directions were not seen. It appears neurons in layer fc4 do not have specific selectivity to translational motion direction; they instead respond to all translational directions more or less equally.

**Figure 15:**
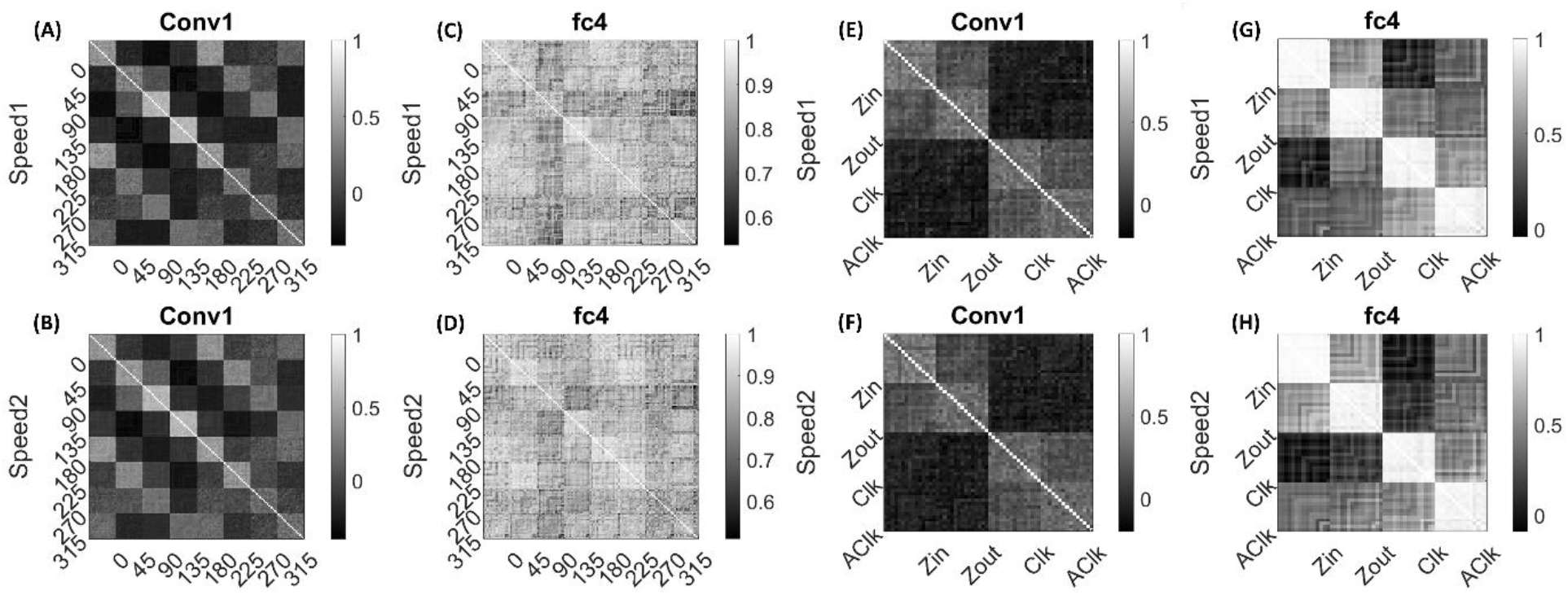
Correlating Conv1 responses with the cell responses in motion hierarchy: Plots display RSMs for CNN layer (conv1) and last hidden layer (fc4). Matrix entries in (A) – (D) indicate the pairwise correlation coefficients calculated for the translational responses, separately for each speed. Matrix entries in (E) - (H) indicate the pairwise correlation coefficients for the optic flow motion responses, separately for each speed. Elements of the matrices are grouped according to the direction and flow type as indicated along the axes.

Figs. 15E and F display RSMs corresponding to optic flow stimuli in the conv1 layer, showing clear distinction between radial and circular motion types, with less prominent selectivity to specific flow type. Whereas, as shown in Figs. 15G and H, very strong block-diagonal along with clearly distinct RSM entries of fc4 layer, indicate that the units are highly selective to different flow types. To understand more about the above conv1 and fc4 responses and their selectivity to various types of translational and flow sequences, we built and trained a linear classifier. In case-1where perceptron trained with ‘conv1 responses to translational stimuli’ produced good recognition accuracy on the test set in both the cases: 85% in speed-1 and 87% in speed-2. In case-2 where perceptron trained with ‘fc4 responses to translational stimuli’ produced less recognition accuracy on the test set in both the cases: 37% in speed-1 and 48% in speed-2. In case-3 where perceptron trained with ‘fc4 responses to optic flow stimuli’ produced high accuracy on the test set (100% in both cases), However, in Case-4 where perceptron is trained on ‘conv1 responses to optic flow stimuli’ produced high accuracy (95% in both cases). It appears that lower layers in the CNN develop selectivity to translational motion while the higher layers code for only the optic flow motion. In sum, bottom-to-top layers of model-3 gradually shifted from direction selectivity of V1 cells to local flow motion estimation and finally optic flow type selectivity, which is reasonably consistent with the idea of functional hierarchy in the macaque motion processing (Born & Bradley, 2005; Duffy & Wurtz, 1991a, 1991b; Hubel, 1995).

## 4. DISCUSSION

Over the last decades, vision research had unraveled a cascade of motion processing stages within the hierarchy of visual cortical areas (V1-MT-MSTd) along the dorsal pathway. V1 neurons have small receptive fields (∼0.5° – 2°) (Duffy & Hubel, 2007; Zeki, 1993) and therefore can analyze movement over only a tiny portion of the visual field. The extra-striate middle temporal area (MT or V5) has a receptive field 10 times the size of V1, while still covering only a relatively small fraction (∼2° - 15°) (Komatsu & Wurtz, 1988a; Richert et al., 2013; Saito et al., 1986) of the visual field. However optic flow covers the entire visual field. The medial superior temporal area (MSTd) has receptive fields (>30°) (Amano et al., 2009; Raiguel et al., 1997) that cover large parts of the visual field (Komatsu & Wurtz, 1988b; Saito et al., 1986) are said to be coding for optic flow motion. The three models described in this paper were based on this functional hierarchy and learn to recognize the type of optic flow present in the given dot sequence.

Here first we list out some of the key features of experimental MSTd cell responses (Duffy & Wurtz, 1991a, 1991b, 1995; Graziano et al., 1994; Graziano, 1990), to interpret the performance of the proposed models. (i). The receptive fields of MSTd cells are (> 30°) much larger than those of the MT cells. (ii) MSTd cells respond to different types of motion stimuli such as unidirectional planar/translational motion, clockwise and counter-clockwise rotational motion, outward and inward radial motion, and various spiral motions (Duffy & Wurtz, 1991a, 1991b; Graziano et al., 1994; Graziano, 1990). (iii). Some MSTd neurons respond maximally to the specific motion type and less strongly, to neighboring motion types (Duffy & Wurtz, 1995) while the others respond moderately to all motion types. There are neurons that do not respond selectively to any of the motion components (Duffy & Wurtz, 1991b; Graziano et al., 1994).

Based on the results provided in Sections 3.1 and 3.2 we can summarize the response properties of neurons in model-1 and model-2 as follows. (i) The neurons in the OFNW simulating MSTd neurons have much larger receptive fields than the neurons in DSMN. Also, they respond to different types of optic flow motion stimuli, including radial and circular motions. (ii) Many neurons in CPNW and HBNW respond most strongly to their preferred translational motion direction, while they also respond less strongly to neighboring directions and moderately to opposite directions. These features match well with the properties of experimental MSTd neurons.

However, neurons in MT that respond to different translational motion directions form a continuum of response selectivities instead of discrete classes (Recanzone et al., 1997) similarly the case with MSTd neurons. In CPNW, since there is no competition across cell-planes, neurons form a discrete class with different directional preferences (i.e., all the neurons in one cell-plane own same direction preference and neurons in different cell-planes exhibit different direction preferences), which is not in agreement with properties of neurons found experimentally (Duffy & Wurtz, 1991b; Graziano et al., 1994; Recanzone et al., 1997). We developed Hebbian network (HBNW) by incorporating competition across the neurons as described in Section 2.4. Through competitive learning, HBNW neurons along each vertical column are trained to respond selectively to the different motion directions, which form continuum of response selectivity along 2D array of neurons. This continuum of responses divided, relatively evenly and randomly, into a set of clusters, each represented by a particular output neuron in OFNW, which is consistent with the empirical findings (Duffy & Wurtz, 1991b; Graziano et al., 1994; Recanzone et al., 1997). Thus, for the presentation of each class of translational motion sequence, the two-stage network (DSMN + HBNW) exhibits a sparcely distributed set of active neurons which represents that translation direction the feature space, which is typical for competitive learning.

Also, HBNW winners corresponding to each input flow sequence are arranged in accordance with the pattern of that optic flow type. In other words, as shown in Fig. 10, ‘HBNW Resp’ displays radial arrangement (winner preferences) for the presentation of radial motion. Similarly, a circular arrangement for the presentation of rotational motion. This is consistent with the hypothesis proposed by various researchers (Saito et al., 1986; Tanaka & Saito, 1989) that the receptive field of an MST cell responsive to circular or radial motions is composed of a set of directionally selective MT cells arranged in accordance with the pattern of that optic flow component.

The first two models presented in this paper simulate direction selective properties of V1 cells, the local flow selective responses of MT cells and selective responses to various optic-flow motion types found in the MSTd area. Both these models show that sophisticated neuronal responses to motion stimuli can be accounted for by relatively simple network models. On the other hand, recent deep convolution neural networks (CNNs) have emphasized layer-wise quantitative similarity between convolutional neural networks (CNNs) and the primate visual ventral stream (Yamins & DiCarlo, 2016). However, whether such similarity holds for the motion selective areas in motion pathway, is not clear through above studies.

In the studies with model-3, we investigate whether CNNs can reproduce the tuning properties observed in the visual motion areas of the macaque brain. We explore the correspondence between the trained model-3 CNN layers and the macaque motion areas by calculating RSMs. Note that we did not constrain model-3 to match neural data, instead by comparing RSMs corresponding to translational and flow motion sequences at convolution layer (conv1) and last hidden layer (fc4), we showed that top output layer is highly predictive of MSTd responses and the intermediate convolutional layer (conv1) is highly predictive of neural responses in MT, an intermediate motion area that provides the dominant cortical input to MSTd. The correlation results show that, in model-3, as one traverses from the input to the output layer, response selectivity gradually shifts from direction selectivity (in VSMN), to local flow selectivity (in conv1), to flow type selectivity (in fc4), which is consistent with the idea of functional hierarchy in the macaque motion pathway. Furthermore, these studies indicate that CNN, in combination with basic sequence processing capabilities offered by DSMN, can be used to build quantitative models of motion processing.

### 4.1 Biological relevance of Model-2

Neurons in the individual NFs of DSMN are designed to have center-surround lateral connectivity, where the lateral connections are trained by asymmetric Hebbian learning. All these centers surround lateral connections and afferent connections are adapted through asymmetric Hebbian rule. In asymmetric Hebbian learning rule, the correlation between the presynaptic state at the current time, and the postsynaptic state at a later time is used. Correspondingly, while training HBNW the symmetric Hebbian rule is used (Hebb, 1949, 2005) wherein the pre- and postsynaptic states are considered at the same instant. If we combine this with the winner take all rule, post synaptic neurons compete with each other and the neurons that produce the largest response becomes the winner. Only the winner’s afferent connections are adapted by Hebbian learning. The winner-takes-all rule used in HBNW facilitate the competition cross the neurons so that different neurons become selective to the different motion directions present in the input. The adaptable lateral connections, Hebbian rule and winner take all mechanisms are biologically plausible and have been tested experimentally (Salzman & Newsome, 1994). Likewise, the initial afferent response of the neurons in DSMN are passed through the piecewise-linear sigmoid function and in HBNW the responses are passed through sigmoid function to make the response nonlinear. Experimental studies have shown that responses of cells in visual cortical areas show significant nonlinearities depending on spatiotemporal activity distribution and also such response nonlinearities have been demonstrated in the LGN and in area V1 and beyond (Solomon et al., 2010; Williams & Shapley, 2007).

### 4.2 Comparison between model-2 and model-3

Model-2 follows a bottom-up approach where neurons in the lower stages are trained first followed by the higher stages in the hierarchy. The initial/first stage neurons are trained to recognize the direction of moving stimuli present within the receptive field; the middle stage neurons are trained with translational dot sequences to encode the direction of local flow; and the last/output stage neurons are trained using different optic flow type sequences. In each stage, training is designed based on the experimental response properties of the neurons present at various levels of motion pathway. This manner of modeling requires not only the knowledge of neurobiological findings but also a good grasp on various neural modeling approaches.

On the other hand, deep learning models are completely data driven, easy to design and train, and require very little pre-programming and domain knowledge. Moreover, various studies (Kriegeskorte, 2015b; Kriegeskorte et al., 2008; Kriegeskorte & Kievit, 2013; Mur et al., 2013; Yamins et al., 2013) demonstrated the parallels along the hierarchy between layers of CNN and the visual areas of ventral pathway. Model-3 is trained end-to-end directly using optic flow motion sequences without any human intervention. Simulation results showed that CNNs can explain the experimentally observed tuning properties of motion areas MT and MSTd and also exhibit the representational similarity with motion areas.

## 5. CONCLUSION

In this paper we simulated three models to recognize the type of optic flow present in the input sequence. All the models explain different functional properties of neurons present in the motion pathway. Further model-3 can be viewed as the candidate models to explain the different aspects of motion processing apart from optic flow. In the future, we further would like to simulate model-3 to understand other motion aspects such as structure from motion and recognition of biological motion.

## AUTHOR CONTRIBUTIONS

AG performed designing, coding, running simulations and manuscript preparation. VC performed designing the model and manuscript preparation.

## Footnotes

1 however, more recent studies have identified direction sensitive cells as early as in the retina (Wei et al., 2011; Wyatt & Daw, 1975)

